# A genome-wide, machine learning-guided exploration of the *cis*-regulatory code involved in neuronal differentiation

**DOI:** 10.1101/2025.05.14.653995

**Authors:** Océane Cassan, Julien Raynal, Christophe Vroland, Kayoko Yasuzawa, Tsukasa Kouno, Jen-Chien Chang, Chung-Chau Hon, Jay W. Shin, Masaki Kato, Hazuki Takahashi, Takeya Kasukawa, Tomoe Nobusada, Vincenzo Lagani, Robert Lehmann, Kévin Yauy, Piero Carninci, Chi Wai Yip, Laurent Bréhélin, Charles-Henri Lecellier

**Affiliations:** LIRMM., Univ Montpellier, CNRS, Montpellier, 34000, France.; Institut de Génétique Moléculaire de Montpellier, University of Montpellier, CNRS, Montpellier, 34000, France.; RIKEN, Center for Integrative Medical Sciences, Yokohama, 610101, Kanagawa, Japan.; Biological and Environmental Science and Engineering Division, King Abdullah University of Science and Technology (KAUST), Thuwal, Saudi Arabia.; Institute of Chemical Biology, Ilia State University, Tbilisi, Georgia.; Département de Génétique Médicale, Maladies Rares et Médecine Personnalisée, CHU Montpellier, Montpellier, France.; Human technopole, Human technopole, via Rita Levi Montalcini 1 Milan, Italy.

**Keywords:** Guided clustering, Interpretable Machine Learning, Neuron differentiation, Single-cell ATAC-seq, Single-cell RNA-seq, *Cis*-regulatory code, Regulatory genomics

## Abstract

Gene expression is controlled by proximal and distal *cis*-regulatory elements (CREs), containing DNA motifs bound by various transcription factors (TFs). Other sequence features, such as specific k-mers or low complexity regions, have also been implicated [1–3]. However, in a dynamic biological process such as cell differentiation, we lack an understanding of how the transcriptional activity of CREs progressively change and what sequence features underlie these transitions, which may reflect common and/or coordinated regulatory processes. Here, we use single-cell ATAC-seq and RNA-seq to follow, at a genome scale, CREs along differentiation of induced pluripotent stem cells into cortical neurons and develop a method to automatically identify the diversity of CRE profiles and their underlying sequence features. We propose a machine-learning guided clustering algorithm, STOIC (Statistical learning TO Inform Clustering), that jointly learns an unsupervised clustering of the CREs in the space of the activity profiles and a supervised predictor associated with each cluster in the DNA-sequence space. STOIC is specifically designed to provide readily interpretable results. We show that the method identifies CRE profiles associated with highly predictive sequence features and outperforms methods solely concerned with co-activity clustering on this task. Orthogonal data collected in the same settings link the inferred CRE clusters to specific enhancer or promoter signatures. Furthermore, we show that the DNA features unveiled by STOIC reflect biologically relevant regulators and offer a valuable basis to dissect elements of the cisregulatory grammar. Finally, we demonstrate the general applicability of STOIC by analyzing five bulk CAGE datasets of human cells responding to various treatments.

## Main

The gene expression program is controlled by DNA regions called *cis*-regulatory elements (CREs), which are either proximal (promoters) or distal (enhancers) to gene start sites, and whose activity must be tightly orchestrated to ensure proper cell biology and functions. To decipher the *cis*-regulatory code [4], it is therefore crucial to understand how regulatory information is read in the genome, i.e. how and when CREs exert their action. Current efforts indicate that this problem can be solved to a large extent at the level of the DNA sequence. CREs harbour specific sequence features (motifs, k-mers or low complexity regions [1, 3, 5–7]), that are recognised by *trans*-regulatory proteins, typically transcription factors (TFs), and that may be specific or common to enhancers and promoters [8, 9]. CRE activity is linked to chromatin opening, which can be measured by ATAC-seq (*Assay for Transposase-Accessible Chromatin with high-throughput sequencing* ) [10]. CREs are also sites of active transcription, which can be measured by different RNA-sequencing protocols, including 5’end RNA-seq [11]. While transcription is traditionally used to assess promoter activity, it is also thought to correlate with enhancer activity [12–15]. Chromatin opening and transcription are therefore two hallmarks of CRE activity. Excitingly, it has become possible to probe CRE activities during a dynamic process such as cell differentiation or in response to a given stimulus (pathogen, drug, . . . ) at the single cell level with single-nucleus ATAC-seq and single-cell RNA-seq [16]. This provides valuable resources to finely identify which CREs are working in a coordinated manner and what is their underlying regulatory DNA grammar.

Understanding how chromatin and transcription patterns are determined, *i.e.* how the activity of CREs progressively changes over time as a function of their sequence has been a long-standing problem and the focus of many studies. Several types of approach have been proposed. First, CRE activity in a given time point or cell type can be linked to the presence of specific sequence features. To do so, some methods embed cells into the space of the DNA sequence features found in their regions of open chromatin, like chromVar and BROCKMAN [17, 18], or by embedding cells into a joint accessibility and k-mers space as in CellSpace [19]. However, these methods infer transcriptional regulatory programs by partitioning cells, rather than directly CREs, thereby identifying a mixture of CREs per cell type whose activities are not necessarily coordinated in other cell types and/or along the dynamic cellular process under consideration. Moreover, these methods lack an objective metric to directly assess the association between sequence and activity. Indeed, the identified features, although truly enriched in certain cell clusters, may not be linked to CRE activity.

A second type of approaches thus chose a supervised setting, where a machine learning (ML) model is trained to predict CRE activity from the DNA sequence. In this framework, the accuracy of the model provides a useful and objective measure of the association between sequence and activity. Several approaches have been proposed for this—whether activity was measured by chromatin opening or transcription level, from linear and logistic models to random forests (RF) or deep learning approaches based on convolutional neural networks (CNNs) [3, 20–25]. With these approaches, a ML model that predicts CRE activity level on the basis of DNA sequence is learned on each time point (either independently or simultaneously using a multi-task approach). For example, scover [20], an innovative supervised strategy tailored for single-cell datasets, predicts CRE activity in homogeneous groups of cells using DNA sequence. Sequence features important for the predictions can then be extracted from each model. Interpretation strategies vary between models that directly take predictive features as input such as random forests (RFs) and linear models [26–28] or models based on raw DNA sequence and performing *de-novo* feature extraction via CNNs [28–31]. However, while these supervised methods can associate sequence features to each time points or cells, they are not sufficient to identify regulatory elements associated with entire and coordinated CREs activity profiles. For this, sequence features importance in each time points or cells should be compared: this contrasting post-processing and interpretation of a potentially large number of models may neither be trivial nor efficient, and, to the best of our knowledge, appropriate approaches are lacking.

Lastly, in a third type of approaches, CREs are clustered based on their activity profile in an unsupervised manner [32–35], and then specific sequence features are associated with each CRE cluster using motifs discovery tools [36]. One drawback is that, unlike supervised approaches, this treats each motif individually and does not build a multivariate model of CRE regulation from DNA sequence. Furthermore, the results of these two-steps approaches are highly dependent on the chosen clustering method and parameters (especially the number of clusters), that may yield drastically different results in terms of motif discovery. Indeed, in a given clustering, the frontiers between clusters are often blurred or uncertain, and other clustering solutions can be equally likely. As a consequence, the inferred clustering does not guarantee that CREs belonging to the same cluster actually share common sequence features, nor that the clusters correspond to regions of the activity space where sequence features enrichment is maximal, making it hard to link DNA sequence with activity [34].

Ideally, we would like to have a method that combines the supervised element of the second category of models, which provides an objective metric to assess the associations between sequence features and activity, together with the ability to directly learn the sequence features associated with entire activity profiles of the third category. Therefore, we propose a method called STOIC (Statistical learning TO Inform Clustering) to co-optimize the clustering of CREs and the sequence features discovery using an interpretable machine learning approach, allowing the discovery of regulatory motifs and their contribution to gene expression patterns simultaneously. Importantly, STOIC is based on a supervised approach that allows to objectively measure the global strength of the association between DNA sequence and CRE activity but also the specific importance of each DNA feature in the predictions. With this new approach, we leverage combined single-nucleus ATAC-seq and single-cell RNA-seq data to characterize the DNA grammar orchestrating human neuronal differentiation. We show that the CRE profiles identified by STOIC are associated with strong sequence features and that many of these sequence-to-activity associations cannot be identified when CRE clustering and sequence features discovery are done separately, without co-optimization. The interpretation of STOIC clusters reveal typical enhancer or promoter signatures, and the sequence features extracted from their supervised models unveil important TFs and DNA patterns for neuronal differentiation.

In order to assess the general applicability of STOIC to datasets of diverse nature and dimension, we also applied STOIC to bulk CAGE (*Cap Analysis of Gene Expression*) data from various kinetic experiments [13]. In summary, our approach automatically explores the complexity of the *cis*-regulatory code at genome scale and provides an updated perspective on the transcriptional regulations at play during several biological dynamic processes. STOIC, together with notebooks, are provided as an R package available in the repository https://gite.lirmm.fr/ml4reggen/stoic.

## Results

### STOIC is a ML approach for the clustering of CRE activity profiles guided by DNA sequence

The principle of STOIC is to link regulatory DNA features to coordinated CREs activity profiles. To do so, STOIC uses an original approach that performs a non-supervised clustering of the CREs in the activity space, using a supervised predictive model in the sequence space as a guide (Fig. 1). Briefly, STOIC delineates clusters of co-active CREs, and the parameters of each co-activity cluster—*i.e.* their centroid and size (radius) in the activity profile space—are optimized in order to maximize the predictive accuracy of a supervised model associated with the cluster. This supervised model is a random forest (RF) [37] trained to predict whether a CRE belongs to the cluster or not, based solely on its DNA sequence. In this study, sequences were described by the scores achieved by a set of transcription factor binding motifs (TFBMs) from the JAS-PAR 2024 database [38] and by the frequency of k-mers in the sequence (see Methods). By iteratively fitting the cluster parameters and the RF in order to improve the accuracy of the predictions, the cluster progressively converges to an area of the activity space with stronger association with sequence information. Clusters are sequentially optimized in this way until the entire activity space has been explored, and the full procedure is repeated with different random initializations to improve stability. Note that STOIC does not necessarily assign all CREs to a cluster, leaving aside regions of the activity space where no link with DNA sequence could be captured by supervised models. The output of STOIC is a collection of non-redundant, optimized co-activity clusters and their associated supervised model (RF), from which the most important sequence features can be extracted. The detailed algorithm of STOIC can be found in the Methods.

**Fig. 1.**
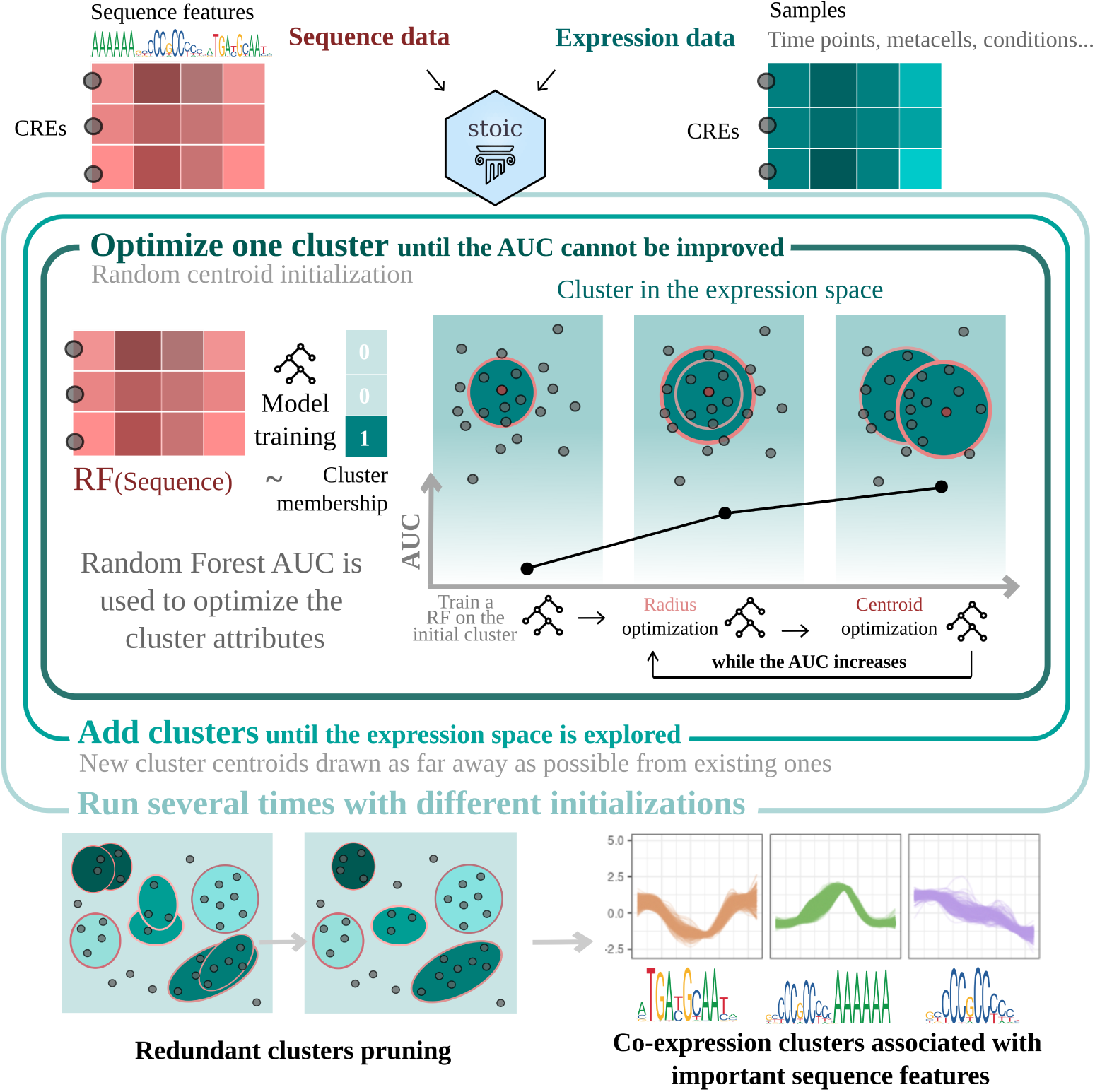
STOIC carries out the co-activity clustering of CREs guided by their sequence features. A cluster is defined by two parameters: its centroid and radius in the activity profile space. For each cluster, a supervised model is trained to predict whether a CRE belongs to the cluster using only sequence features as predictors. The AUC of this model is then maximized by iteratively updating the cluster parameters until convergence. This is repeated for different clusters until the activity space is fully explored, and then repeated for different random initializations. Redundant clusters are then pruned based on model swaps, correlation and overlap. STOIC’s output is a set of clusters, each of them associated with a supervised model that can be interrogated to learn the most important cluster-specific sequence features.

### Neuronal differentiation is studied based on single-cell RNA-seq and single-nuclei ATAC-seq

We propose to apply STOIC to study the regulatory grammar underlying human neuronal differentiation. For this, we rely on a single-cell 5’-end RNA-seq (scRNA-seq) and single-nuclei ATAC-seq (snATAC-seq) dataset of induced pluripotent stem cells (iPSCs) differentiating into neural stem cells (NSCs) and finally cortical neurons (Fig. 2a, Methods). SEACells [39] was used to group cells into 391 metacells according to the RNA and ATAC modalities. These 391 metacells, *i.e.* homogeneous cell clusters corresponding to granular cell types, were ordered along the differentiation pseudotime (Methods, Table S1). The CREs of this study are genomic regions located under peaks of snATAC-seq. The activity of 10912 CREs was measured by the scRNA-seq signal within these regions in the different metacells (Fig. 2b, Methods, Table S2). In the rest of this article, CRE activity is therefore synonymous with the expression level of open CREs. These CREs can have various activity profiles and peak at different moments of differentiation (Fig. 2c-d).

**Fig. 2.**
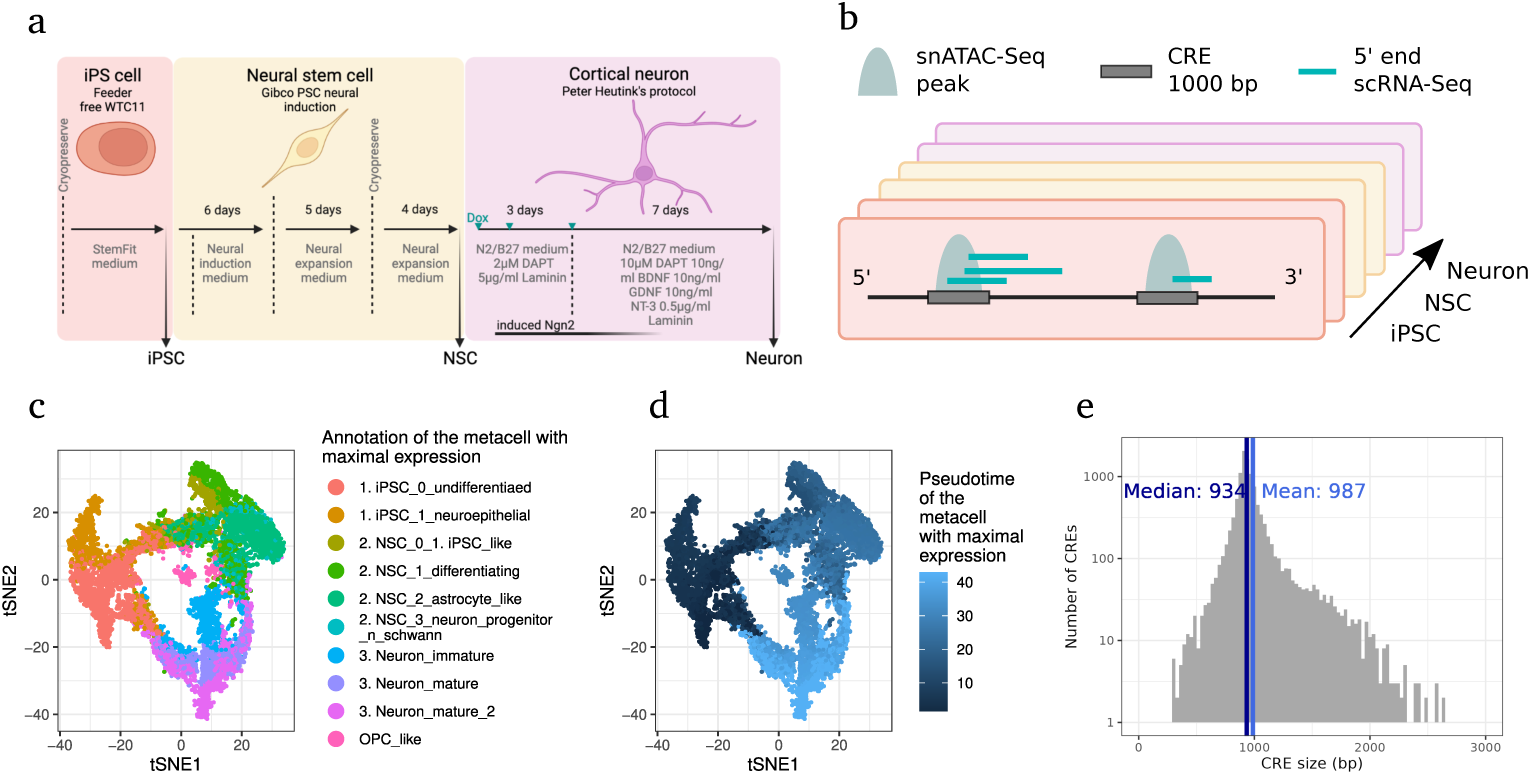
Single-cell neuronal differentiation dataset. **a.** Protocol used to differentiate iPSCs into cortical neurons. **b.** Schematic view of the CRE sequences, located under single-nuclei ATAC-seq peaks, and their transcriptional activity provided by single-cell 5’-end RNA-seq. Single cells were aggregated into 391 metacells, in which the ATAC-seq and RNA-seq signals were pooled. **c-d**. Representation of transcriptionally variable CREs in the expression space. The 2-dimensional expression space was obtained from the original 391-dimensional space via a tSNE. Each dot represents a CRE, colored depending on the metacell annotation where it reaches its maximal expression (c), or depending on the pseudotime at which it reaches its maximal expression (d). **e.** Size distribution of the CREs ATAC-seq peaks used to define CREs.

In order to extract DNA features from this CREs and analyze them with STOIC, we first had to define the length of the CRE sequences to consider. We tested sequences of length 100bp, 200bp, 500bp, 1000bp and 2000bp centered around snATAC-seq peak mid-positions (Fig. 3a). From these sequences, we extracted k-mers frequencies (44 mono, di and tri-nucleotides) and the TFBM scores of 382 expressed TFs (Methods). The AUC distribution of STOIC’s models was maximal with sequences of length 500 and 1000 bp. At 100 or 200 bp, the AUC decreases, as some predictive sequence features are probably missed by the small DNA window. At 2000 bp, the AUC is slightly lower than at 1000 bp, suggesting that no additional information is learned on larger windows. We finally chose a CRE length of 1000 bp for the rest of this study, also because it is closer to the observed ATAC-seq peaks length (Fig. 2e, Tables S3 and S4).

**Fig. 3.**
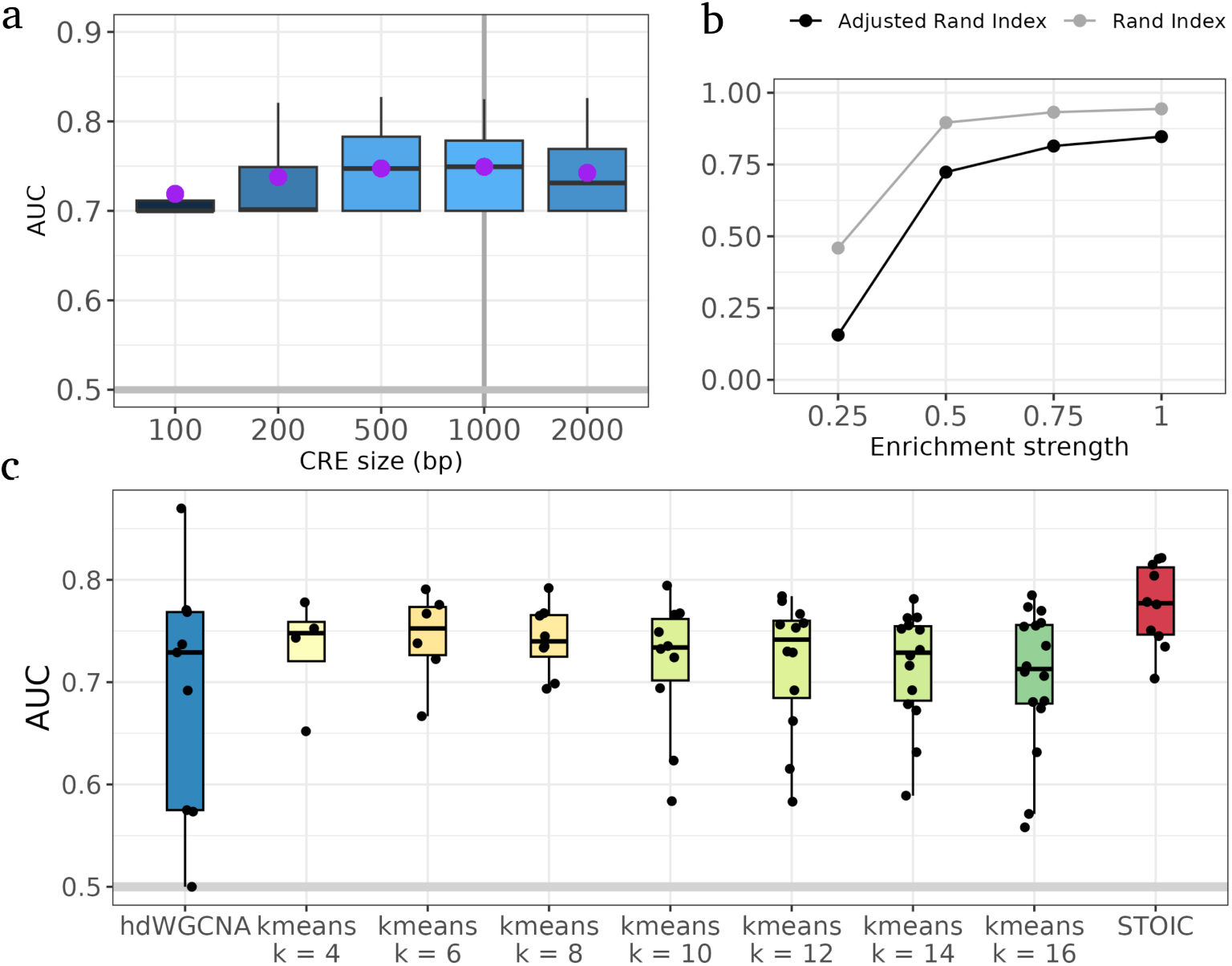
STOIC’s accurately retrieves coordinated activity profiles with characteristic sequence features. **a.** AUC distribution of CREs as a function of CRE sequence size. To compute these AUC distributions, each CRE receives the AUC of its cluster, or 0.7 (the AUC threshold for a cluster inclusion in STOIC) if it is not in any cluster. The mean AUC is shown by the purple dot. The vertical line at 1000 bp represents the CRE size chosen for this study. **b.** Similarity between the simulated gold standard clustering and the output of STOIC applied to the simulated datasets. Similarity to the true clustering is measured by the Rand Index (grey line) and the adjusted Rand Index (black line), as a function of the strength of enrichment of sequence features in the simulated datasets. **c.** Association strength between sequence features and co-activity clusters in STOIC versus other clustering methods, as given by the AUC obtained when predicting cluster membership using sequence features. Each dot corresponds to a cluster. Methods are ordered by mean AUC. Alternative approaches, *i.e.* HdWGCNA and all k-means clusterings are adjusted to the same coverage as STOIC.

### STOIC recovers highly predictive sequence features and outperforms the clustering approaches on this task

We first ran simulations to assess the accuracy of STOIC in retrieving co-actvity clusters and their DNA features under different levels of noise. To generate a gold standard dataset, we performed a clustering of activity profiles using the true activity data, and then artificially enriched the sequences of certain clusters with one or two specific sequence features, using different enrichment strengths (Methods, Fig. S1a). We observed that, given the simulated sequence features and the original expression data, STOIC is able to recover the clusters of the gold standard, with an accuracy that increases with the enrichment of sequence features (Fig. 3b, Fig. S1c-g).

In addition, for all inferred clusters, the most important feature(s) always correspond(s) to the artificially enriched one(s) (Fig. S2).

We then asked whether the sequence features identified by STOIC were more relevant than sequence features identified by a standard two-step approach that would first cluster CREs on the basis of their activity profiles and then identify the important DNA features associated with each cluster. As alternative co-activity clustering methods, we included the k-means algorithm with different numbers of clusters, and hdWGCNA [33], a weighted correlation method that extracts co-activity modules from high dimensional transcriptomic data (Methods). For each cluster, a RF was then trained to predict cluster membership using the same sequence features as in STOIC. The association strength between co-activity clusters and sequence features as measured by the AUC is always higher in STOIC (Fig. 3c). This observation holds whether the different clusterings are adjusted to have the same coverage (*i.e.*the same number of CREs are in a cluster) or not (Fig. 3c, Fig. S3). This illustrates that the proposed ML-guided approach produces clusters with stronger DNA features associations as compared to a simpler two-step strategy. This is expected, as STOIC has been purposefully designed to maximize this association, which is not the case of other existing approaches.

### STOIC identifies a wide range of co-activity clusters associated with specific sequence features and biologically relevant epigenetic signatures

The application of STOIC to the neuronal differentiation dataset resulted in a collection of 10 CRE co-activity clusters, with varying sizes and activity profiles peaking at distinct developmental stages (Fig. 4a and d, Table S5). All final clusters display AUCs greater than 0.7. The sequence features identified by STOIC revealed a mixture of important TFBMs and k-mers frequencies associated with each cluster (Fig. 4c, Table S6).

**Fig. 4.**
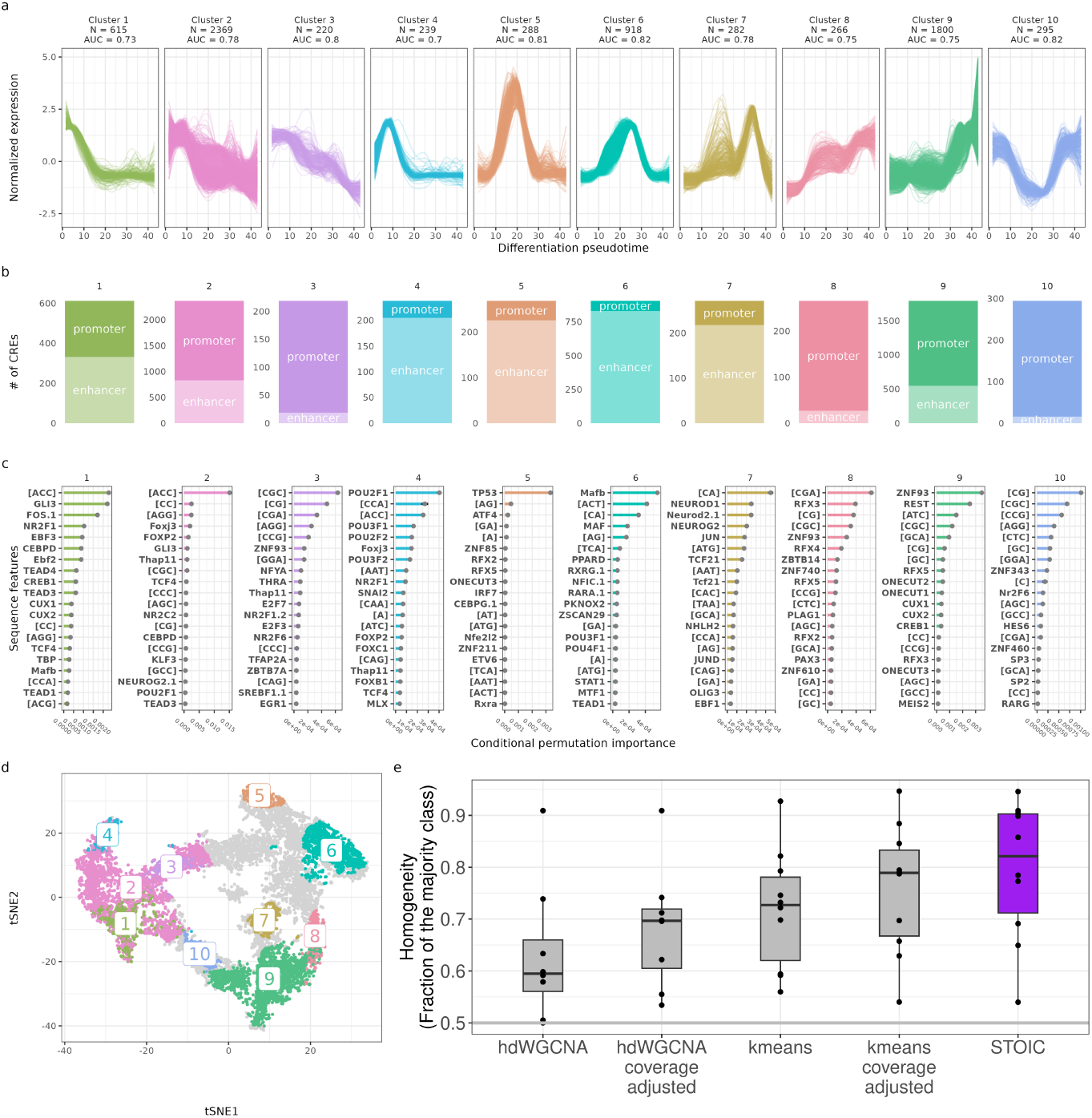
STOIC recovers diverse co-activity clusters with sequence features specific to neuronal differentiation stages and epigenetic signatures. **a.** Expression profile of each cluster (normalized and smoothed) along the differentiation pseudotime. Clusters are ordered by pseudotime of maximal activity. The number of CREs and AUC of the cluster are displayed above each profile. **b.** Enhancer-promoter composition of each cluster as inferred from epigenetic marks (CUT&Tag data). **c.** The 20 most important sequence features in each cluster. Feature importance is extracted from RF models via conditional permutations (Methods). Features within square brackets are k-mer frequencies computed in the entire 1000bp sequence, and features without brackets are TFBM scores. **d.** Representation of STOIC clusters in the reduced expression space (tSNE). Each dot is a CRE, colored by its cluster membership. Grey CREs do not belong to any cluster. **e.** Enhancer-promoter homogeneity of STOIC clusters versus alternative co-activity clustering approaches (adjusted or not to the same coverage as STOIC). Homogeneity is computed as the fraction of the greatest class of CREs, *i.e.*the fraction of enhancers for enhancer-rich clusters and promoters for promoter-rich clusters.

The annotation of CREs based on their epigenetic signatures (Methods, Table S7) showed that most clusters preferentially contain one type of CRE (Fig. 4b): promoters (e.g clusters 3, 8 and 10) or enhancers (e.g. clusters 4, 5, 6 and 7), while a smaller number of clusters contain both in similar amounts (e.g. clusters 1, 2 and 9). Compared to alternative clustering methods, the clusters inferred by STOIC display a stronger homogeneity in terms of enhancers and promoters (Fig. 4e). Indeed, although co-activity clustering alone tends to group CREs with similar enhancers or promoters signatures, this is further enhanced in STOIC where sequence features additionally inform the clustering. This is in line with the knowledge that enhancers and promoters have both distinct activity profiles and DNA properties [12]. In particular, promoter-rich clusters in STOIC are characterised by CG-rich k-mers whereas enhancer-rich clusters contain more AT-rich k-mers (Fig. 4c). Additionally, these epigenetic annotations suggest that neuronal differentiation follows a specific genomic program, characterised by a decrease in the expression of enhancers and genes associated with the stem cell state, followed by a transient activation of enhancers and finally the activation of genes associated with neuronal cell state. Such waves of coordinated transcription were also observed in various time courses covering a wide range of mammalian cells and biological stimuli [13].

Since enhancers and promoters are at the core of chromatin interactions, we asked whether the clusters defined by STOIC reflected long-range DNA:DNA and/or DNA:RNA interactions. The DNA:DNA interactions captured through Hi-C in iPSCs, NSCs and neurons revealed that CREs within the same clusters come into contact more often than with CREs from other clusters (Fig. S4a, Methods). This indicates that STOIC models recover groups of CREs physically closer in the nuclear space.

We also observed DNA:RNA interactions within and between clusters (Fig. S4b, Methods), as given by RADICL-Seq data obtained in iPSCs, NSCs and neurons [Pracana et al., in preparation, Lambolez et al., in preparation]. The clusters involved in more DNA:RNA contacts correspond to promoter-rich clusters active in the considered cell type (*i.e.*cluster 3 in iPSCs, or clusters 8 and 9 in NSCs and neurons). While DNA:DNA contact rates between and within clusters appears to be static during differentiation, the DNA:RNA interactions are more cell type specific, with an especially strong variation between iPSCs and NSCs. This tissue-specificity is in line with the reported cell-type specificity of the RADICL-Seq technology [40].

A total of 67% of CREs were associated with one of the 10 clusters. The projection of the inferred clusters in the reduced activity space (Fig. 4d) revealed that STOIC did not identified clusters in particular areas of activity profile, and we sought to understand why. The forced estimation of a RF model in these areas revealed lower AUCs (between approx. 0.5 and 0.65, Fig S5), meaning that STOIC avoids areas where sequence features provide poor predictive power, as intended. Note that the use of *de-novo* motifs discovered by STREME [36] did not improve AUCs in these regions, nor in STOIC clusters (Fig S5c and S6). This indicates that while TFBMs and k-mers are helpful to explain CRE activity profiles, another part of CREs activity may be explained by different DNA features, and/or features located outside of the considered sequences in the present version of STOIC (e.g. distant regulations, post-transcriptional regulators such as microRNAs or RNA-binding proteins).

### STOIC highlights important transcriptional regulators of neuronal processes

The most important TFBMs in the supervised models of STOIC are often from TFs that have already been characterized for their role in neuronal cells fate like for instance NR2F1 [41], GLI3 [42], REST [43], or the TF families NEUROD/G [44], ONECUT [45] and TEAD [46] and RFX [47]. We first assessed whether the importance of TFs in different clusters along differentiation coincides with their expression patterns. Although not all important TFs have an expression profile closely following the activity profile of their cluster, several of the most important TFs are most expressed when the CREs of their cluster peak in activity (Fig S7, Table S8), supporting their importance for this cluster. A good example is cluster 7, where the most important TFs are specifically expressed in immature neurons, following the CREs activity profile of the cluster (Fig. 5a). Cluster 9 also contains many important TFs expressed in neurons simultaneously with their target CREs, such as ONECUT1/2/3, CREB1 or JUND (Fig. 5b). Moreover, the second most important TF of cluster 5, REST, peaks in iPSCs. REST is actually strongly anti-correlated with the activity profile of this cluster (Fig. 5c), which is in agreement with its documented function of transcriptional repressor [43]. We also cross-compared STOIC’s important TFs with the sysNDD database, a resource curating genes involved in neurodevelopmental disorders. The most important TFs of STOIC models are enriched in TFs of this database (Fig. 5d).

**Fig. 5.**
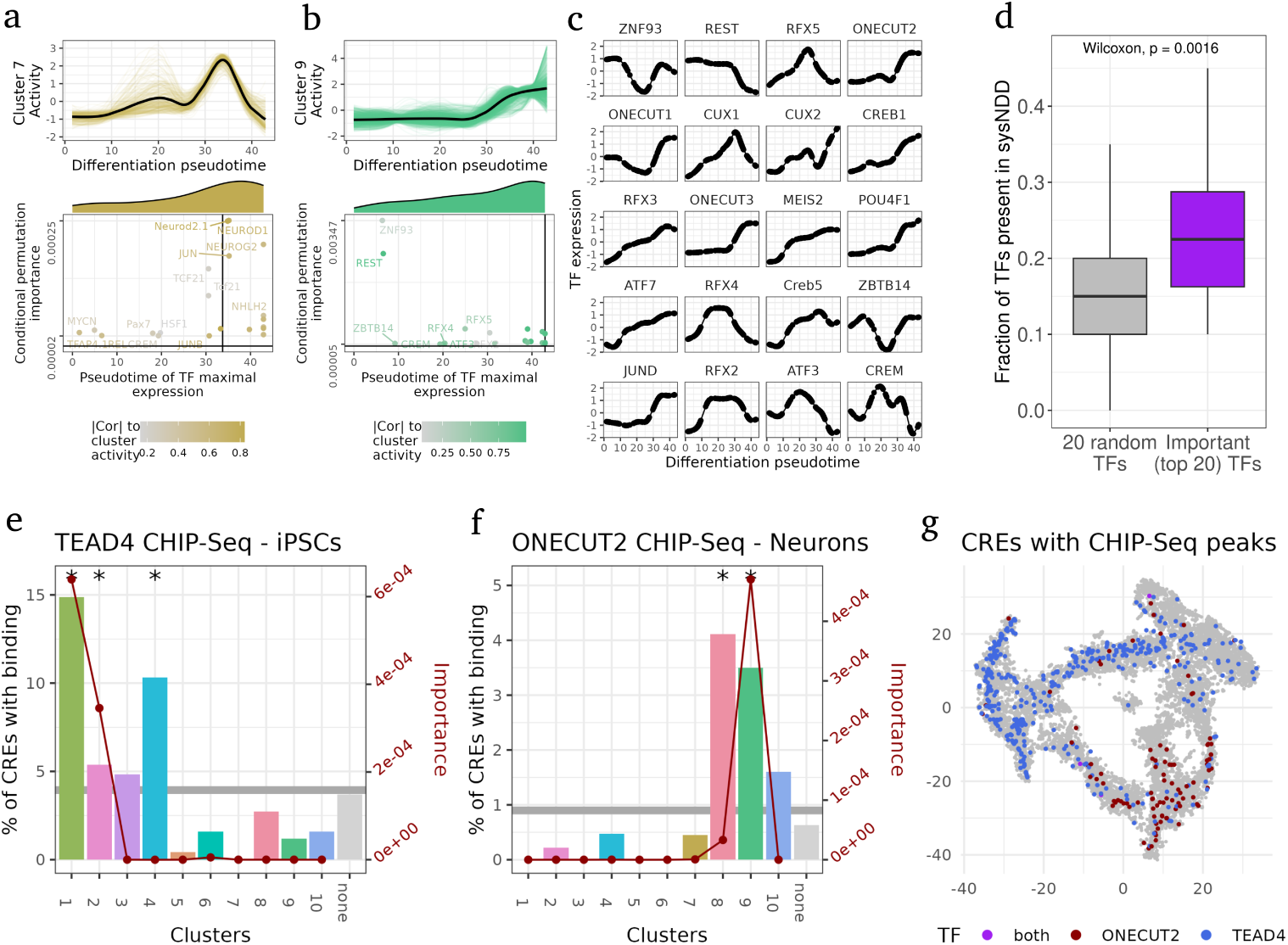
STOIC identifies meaningful transcriptional regulators of neuronal differentiation. **a-b.** Activity profiles of clusters 7 and 9 (top) and, for the 20 most important TFs in these cluster, TF conditional permutation importance as a function of the TF pseudotime of maximal expression (bottom). Important TFs are restricted to the TFs with higher scores in the cluster (*i.e.*enriched and not depleted). The vertical line represents the pseudotime of maximal activity of the co-activity cluster. The marginal distributions of the TFs pseudotimes of maximal expression are shown above the scatter plot. TFs are colored depending on the absolute correlation of their expression profile to the activity profile of the cluster. **c.** Expression profile of the 20 most important TFs in cluster 9. As with the scRNA-seq signal in CREs, TF expression of normalized, smoothed and scaled to a z-score. **d.** Fraction of TFs found in sysNDD in the top 20 TFs of each cluster, versus in the same number of randomly chosen TFs (drawn 1000 times). **e-f.** Percent of CREs in each cluster overlapping a ChIP-Seq peak of TEAD4 in iPSCs (e) and ONECUT2 in Neurons (f). Stars denote a significant (P *<* 0.05) enrichment of binding events based on a one-sided hypergeometric test. The conditional permutation importance of TEAD4 (e) and ONECUT2 (f) is overlaid. **g.** CREs overlapping a ChIP-Seq peak of TEAD4 in iPSCs and ONECUT2 in neurons in the reduced tSNE activity space.

Finally, we used ChIP-Seq to experimentally validate the important sequence features of STOIC, specifically TEAD4 in iPSCs (Table S9), and ONECUT2 in neurons (Table S10). TEAD4 is predicted as an important TF in cluster 1 and 2 where CREs peak in activity during the iPSC stage. The regulatory function of TEAD4 in iPSCs is supported by an enrichment of CREs bound by this TF within these two clusters (Fig. 5e and 5g). Similarly, in cluster 9 (and 8 to a lesser extent), the high importance of ONECUT2 unveiled by STOIC coincides with an enrichment of CREs bound by this TF in neurons (Fig. 5f and 5g). This demonstrates that, in these two CHIP-Seq assays, the important TFs revealed by STOIC are supported by *in-vivo* binding. The fact that some clusters are enriched for the binding of a TF that STOIC did not predict as important (e.g. TEAD4 in cluster 4) may be due to CHIP-Seq capturing the direct binding of a co-factor, and only the indirect binding of the TF. In this case, only the co-factor’s TFBM would be a predictive feature, and not the TFBM of the initial TF. Altogether, these results suggest that the TFs with high importance in the supervised models of STOIC are associated with ChIP-Seq binding, are capable of modelling known activation or repression regulatory relationships, and contain crucial genes for neuronal development and processes.

### STOIC can be used to decipher the *cis*-regulatory grammar underlying neuronal differentiation

In order to better understand the sequence features learned by STOIC, we started by asking whether some clusters shared similar *cis*-regulatory code. To test this, we used the supervised models of each cluster to predict the membership of other clusters. More precisely, in such a model swap, we use the model A (learned on cluster A) to predict CRE membership of cluster B (Fig. 6a). If the AUC achieved by model A on cluster B is lower than that achieved by model B (that was learned on cluster B), this means that the sequence rules learned by the two models are different. If there is no AUC decrease, their sequence features can be considered interchangeable. Results show that few of STOIC’s models are really interchangeable, but we noted that clusters with the same CRE composition (e.g. two promoter-rich clusters or two enhancer-rich clusters) can be more easily swapped than clusters with different enhancer-promoter compositions. For example, clusters 8 and 9 are both promoter rich and their sequence rules are similar. However, not all pairs of clusters with the same enhancer-promoter composition can be swapped: clusters 4 and 7 are both rich in enhancers but their sequence rules are very specific, and this is caused by the fact that they are active at different moments of differentiation (iPSCs, and immature neurons, respectively). In summary, swapping the different supervised models of STOIC clusters revealed that the learned *cis*-regulatory code depends both on the epigenetic status (enhancer versus promoter) of CREs, but also on the developmental stage of activity.

**Fig. 6.**
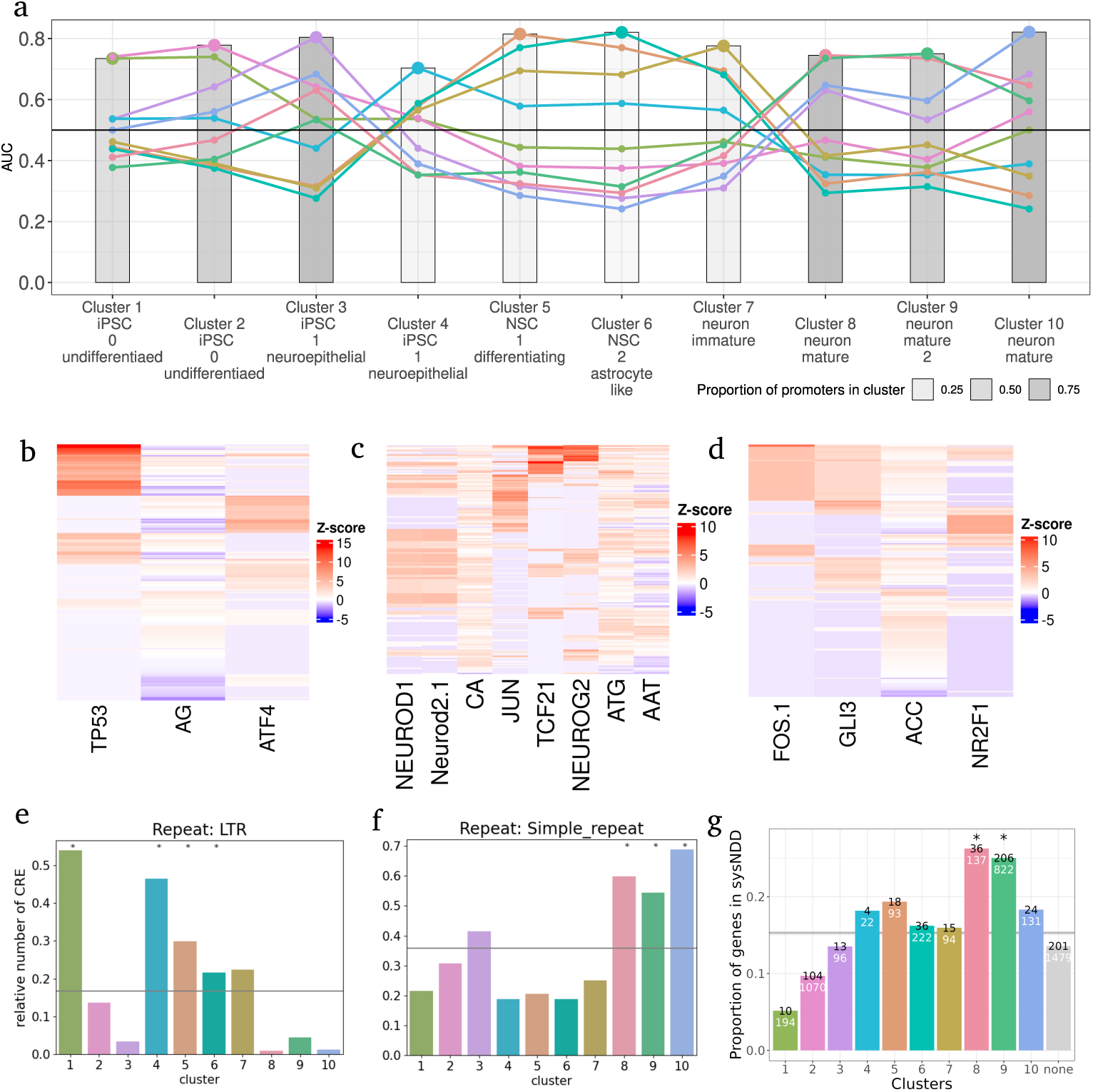
Study of the cis-regulatory code from STOIC models. **a.** Swaps between all pairs of STOIC models. The bars heights are the AUCs of the models applied to predict their own cluster based on CREs sequence features. Each colored line represents the AUC of a given model wen applied to predict the 10 STOIC clusters (its own and the 9 others). The grey scale of the bar color corresponds to the promoter fraction within the cluster. Clusters are ordered and annotated based on the metacells where they reach maximal activity. **b-d.** Heatmap of sequence feature scores of the most important features in clusters 5, 7 and 1 (columns) in the CREs of these clusters (rows). The restricted list of most important features was obtained via the ”elbow” rule from the ranked feature importance curve. Important features were restricted beforehand to the features with higher scores in the cluster than in other CREs. The scores of each DNA feature were scaled to a z-score across the 10912 CREs. Red cells represent scores higher than average, and blue scores below than average. **e-f.** Fraction of CREs in each cluster overlapping LTR repeats (e) and Simple repeats (f) from the RepeatMasker annotation. Stars denote a significant enrichment (P *<* 0.05) of repeats based on a Fisher’s exact test after FDR correction. **g.** Fraction if genes in each cluster contained in the sysNDD database. Stars denote a significant enrichment (P *<* 0.05) based on a one-sided hypergeometric test.

The important k-mers and TFBMs in the models of STOIC were then examined to expand our understanding of the *cis*-regulatory code. For each cluster, STOIC outputs a heatmap of the scores of the most important sequence features in the individual CREs of the cluster (Fig. 6b-d and Fig. S8). Interestingly, many clusters showed patterns of mutual exclusion of TFBMs. For example in cluster 5, TP53 has high binding scores in CREs that are different from the CREs where the second most important TF, ATF4, has its highest binding scores (Fig. 6b). This is also the case for JUN, NEUROD1/2.1 and TCF21 in cluster 7 (Fig. 6c), for which high-score motifs are found in different CREs, or for NR2F1 and GLI3 or FOS.1 in cluster 1 (Fig. 6d). Besides, we pinpoint co-occurrence relationships not only when two TFBMs have extremely similar PPMs like NEUROD1 and NEUROD2.1, but also when the PPMs are different. The latter case could represent co-regulatory relationships like for example TCF21 and NEUROG2 in cluster 7 (Fig. 6c). TFMBs with different PPMs can also co-occur in cases where they are found in CREs originating from the same repeated sequence. This happens for CREs in cluster 1, where FOS.1 and GLI3 have their motifs in similar CREs, that are actually retrotransposable elements (Fig. 6d). This is supported by the strong enrichment of cluster 1 for LTRs (Long Terminal Repeats), a class of repeat elements from the RepeatMasker annotation (Fig. 6e, Table S11). In particular, these LTRs are made of the HERVH-int element flanked by LTR7 elements (Fig. S9a-b, Table S12). This finding in a cluster of CREs specifically active in the iPSC stage is in agreement with previous studies that uncovered the role of retrotransposable elements in the identity and transcriptional regulation of pluripotent stem cells [48]. Along these lines, several pluripotency-related TFs such as OCT4-SOX2 or NANOG are already known to bind LTRs [49]. As the TFBMs identified as important in cluster 1 such as GLI3, FOS.1 or NR2F1 preferentially bind these LTRs (56-63% contain their motif, which is 2-6 folds more frequent than in all CREs), they may also represent novel LTR-binding TFs worthy of future attention.

Finally, STOIC clusters showed enrichments for other types of repeat elements (Tables S11 and S12). For example, SINE and LINE elements were over-represented in enhancer clusters (Fig. S9c-d) in agreement with previous findings revealing their contribution in genomic regulations [50, 51]. Furthermore, simple repeats (or microsatellites) were enriched in clusters of promoters active in neurons, *i.e.* clusters 8, 9 and 10 (Fig. 6f). Upon further inspection, many of these simple repeats turned out to be GC-rich (Fig. S9e-f), which is supported by the importance of GC-rich k-mers frequencies in the models of STOIC (Fig. 4c). Because GC-rich repeat expansions have been linked to 28 genes involved in neuropathologies [52], we intersected these genes with our CRE clusters (Fig. S10). Of the 3 genes associated with GC-repeat expansions and falling in our clusters, one is in cluster 8 (PPP2R28) and two are in cluster 9 (AFF2 and ATXN8OS). Similarly, neurodevelopmental disorder genes from sysNDD are enriched in the same clusters, active in late stages of neuronal differentiation and mature neurons (Fig. 6g). All these elements indicate that the TFBMs, enriched k-mers and low complexity DNA identified on the basis of STOIC have the potential to unveil the grammar of diverse regulatory mechanisms and pathologies associated with neuronal differentiation and neuronal cells.

### Applying STOIC to bulk CAGE kinetic experiments

Finally, we demonstrate that STOIC is not uniquely applicable to single-cell datasets by employing it on several bulk CAGE datasets. These datasets originate from a previous study analysing the transcriptomic reprogramming induced by numerous treatments or differentiation processes in mammalian cells [13]. We chose to revisit 5 of the human datasets in this study, labelled as follows: ARPE (Retinal pigment mesenchyme transition based on ARPE-19 cells), LPS (primary human monocyte-derived macrophages responding to lipopolysaccharide), HRG (MCF7 cells responding to Heregulin), EGF1 (MCF7 cells responding to Epidermal Growth Factor) and IL1b (Primary aortic smooth muscle cells responding to Interleukin-1beta). In these bulk datasets, the number of activity measurements (given by the CAGE signal) per CRE is lesser than in the single cell datasets (between 45 and 75 time points/replicates instead of the 391 metacells). The numbers of transcriptionally variable CREs identified in the original paper are also smaller (between 494 and 7706). The types of sequence features leveraged to describe CREs were the same as in the single-cell dataset (*i.e.*k-mers frequencies and TFBMs of expressed and variable TFs). Overall, the application of STOIC produced several clusters per time course with AUCs ranging from 0.65 up to almost 0.9, and with performances varying slightly between experiments (Fig. 7a). We then focused on two time courses (HRG and IL1b) in order to compare STOIC’s output to the original findings made from these datasets (see Fig. S11 for the detailed results of the three remaining experiments). First, in these two kinetic experiments, we observed that the CRE clusters for which activity is most rapidly induced correspond mainly to enhancer clusters, while clusters with late activation correspond mainly to promoter clusters (Fig. 7b and 7c). This is in accordance with the message of the original study, stating that groups of coordinated enhancers are involved in the earliest transcriptomic response, later followed an activation of promoters [13]. The original authors noted that in the time course HRG, the contribution of the FOS TFBM to transcriptional responses was strong. This was also supported by the results of STOIC, highlighting the TFBMs of FOS (combined to JUN) as important sequence features for a cluster of early-induced enhancers (Fig. 7b). Moreover, in the experiment IL1b, Arner and co-authors identified the high contribution of the nuclear factor

**Fig. 7.**
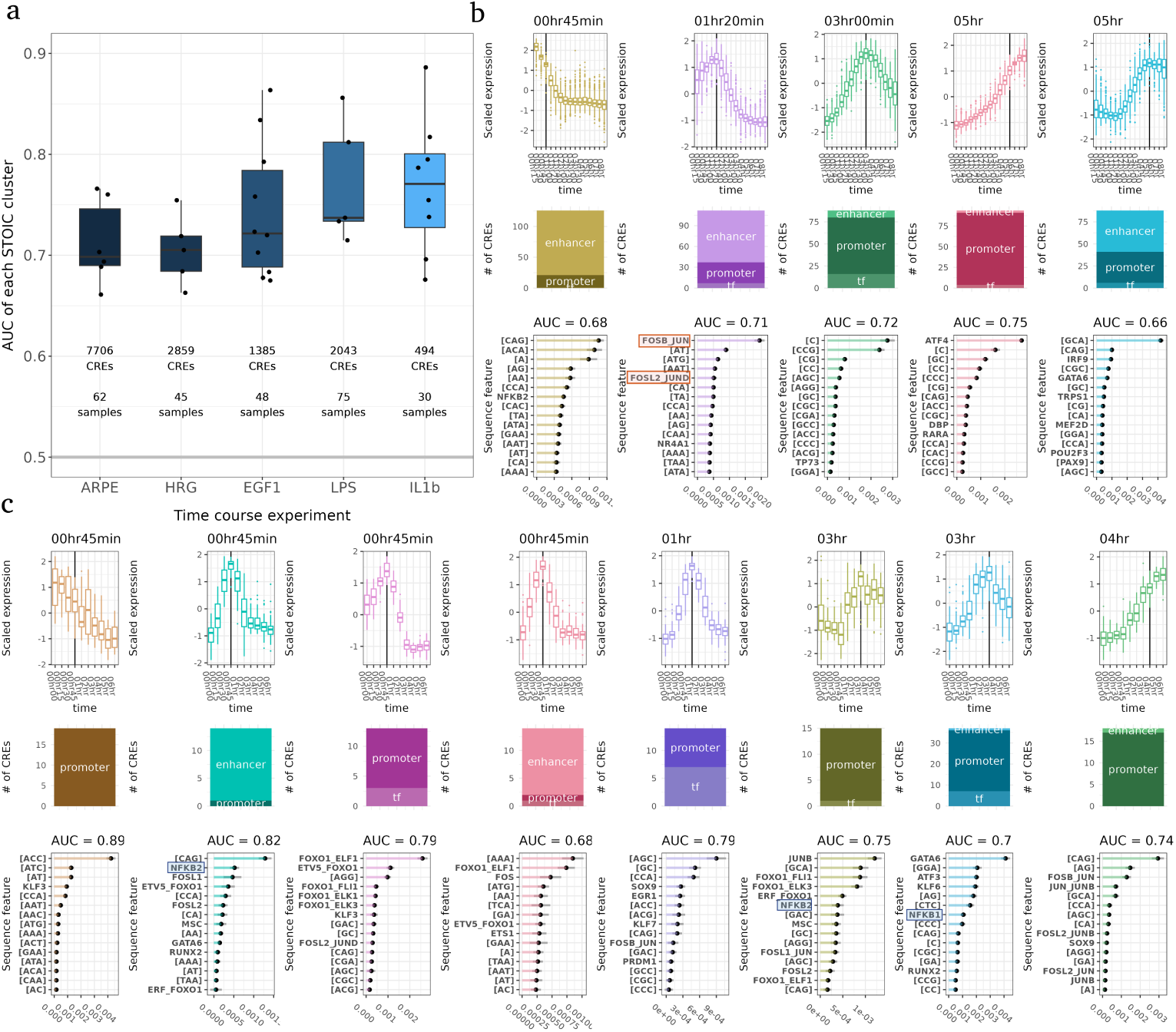
STOIC retrieves coordinated waves of enhancer and promoter activity in bulk CAGE kinetic experiments from [13], along with underlying sequence features. **a.** AUC of each cluster retrieved by STOIC in the 5 bulk CAGE kinetic experiments. The number of transcriptionally variable CREs and the number of time points (samples) is reported for each experiment. **b-c**. Detailed STOIC results on the HRG experiment (b), and on the EGF1 experiment (c). Detailed results include the normalized activity profiles of each cluster (1st row), the composition of the cluster (2nd row), and the important sequence features and AUC in each cluster (3rd row). The vertical line i the activity profiles corresponds to the profile center of mass, *i.e.* the time point where half of the CRE activity has changed. Clusters are ordered by center of mass. CRE annotation: ”enhancer”, ”tf” (for a TF promoter), ”promoter” (for the promoter of a non-TF gene). Sequence features discussed in the original article [13] are highlighted.

NF-*κ*B motif in both enhancers and promoters activation. Accordingly, STOIC identifies several clusters in which NF-*κ*B1 or NF-*κ*B2 are important DNA features: an early-responsive enhancer cluster and two clusters of promoters induced later (Fig. 7c). In addition to these examples, many more new TFs are predicted as candidate regulators of CRE activity profiles by STOIC in these experiments. This includes ATF4 in the intermediate HGR response or FOXO1, ELF1, and GATA6 in the IL1b time course, along with several k-mers frequencies with potential regulatory roles. These results suggest that STOIC provides valuable sequence descriptors of CREs activity profiles derived from bulk experiments, and has the potential to help revisit and refine previously published datasets.

## Discussion

In this work we described STOIC, a novel algorithm to extract DNA sequence features linked to specific activity profiles in CREs. Delineating groups of co-active CREs based on the discriminating power of their DNA sequence offers a clear objective function (*i.e.*the predictive performance of a supervised model) to help parameterize and guide the co-activity clustering. We demonstrate that STOIC has the ability to provide meaningful insights into the regulatory grammar at play in a single-cell resolution dataset of human neuronal differentiation, but also into non-single-cell datasets like CAGE kinetics. In the analysed neuronal dataset, cell fate seems to be mostly defined by the extinction of the pluripotency state accompanied by successive waves of active enhancers in the NSC stage, followed by the establishment of neuronal promoters and enhancers. STOIC clusters are characterized by specific TFBMs, k-mers enrichments, and repeat elements: the regulatory function of several of them was supported by orthogonal experiments like ChIP-Seq assays, and expert-curated databases of neurodevelopmental pathologies. They provide a valuable basis to further dissect the *cis*-regulatory code.

Nevertheless, several limitations of STOIC should be reminded. The sequence features used in STOIC are limited to the TFBMs from JASPAR and k-mers frequencies. This does not allow to model the TFBMs of TFs absent from motif databases, or other forms of sequence patterns. A practical solution to this problem is to supplement STOIC with TFBMs identified with *ab initio* motif discovery algorithms, although this approach did not enabled the discovery of interesting new motifs in our experiments.

A perspective for future work could be to replace the RFs in STOIC by convolutional neural networks, that can extract useful features directly from DNA sequence, like the deep-learning based approaches mentioned in the introduction [20, 22–25]. However, this would inevitably reduce the interpretability of the approach, as measuring CNN variable importance remains a difficult task. Another part of the unexplained CRE activity in STOIC likely stems from the unknown distal enhancers regulating promoter CREs: the inclusion of spatially close enhancer sequences may increase predictive performance.

Another drawback that STOIC shares with all approaches predicting activity from sequence is the correlation between input features (motifs and k-mers). As a consequence, the predictive model learned from the data may not be the only one ”good” model, an issue coined as the Rashomon effect in statistics [53, 54]. To partly overcome this effect connected to model instability, a strategy is to use ensemble methods, which are less sensitive to this issue because they aggregate the predictions of a very large set of predictors [53]. This is the approach used by STOIC, which is specifically based on RFs. Other approaches that have been proposed involve measuring feature importance with regard to sets of equally accurate models [54], or to decompose the importance of a feature by measuring its contribution in different groups of features [55, 56].

An intriguing observation is the existence of sub-groups of CREs within a cluster characterized by distinct DNA features (Fig. 6a-c, and S8). This challenges the idea that all CREs of a co-activity cluster have the same regulatory grammar, as the RFs in STOIC sometimes rather learned features found in distinct sets of CREs. We may hypothesize that even though these CREs have different sequence rules, they are brought together spatially by chromatin folding and show, *in fine*, similar activity profiles. Alternatively, we may suppose that CREs within a co-activity cluster are targeted by the TFs of different signalling pathways controlling similar CRE activity profiles. Under this scenario, distinct CREs would contain the TFBMs of the different pathways effectors. For instance, TP53 and ATF4 target different CREs in the cluster 5 of STOIC, and have already been documented to regulate common processes through independent converging pathways [57].

## Material and Methods

### Single-cell neuronal differentiation dataset

#### Cell culture, differentiation, and single-cell sequencing

The full cell culture and differentiation protocol is provided in Supplementary Methods B. Briefly, the Human iPSCs line i^3^N is a gift from Dr. Michael Ward from NIH and derived from WTC11 iPSCs by introducing a doxycycline-inducible mNGN2 transgene at the AAVS1 site. NSCs were differentiated and maintained by using the PSC Neural Induction Medium (NIM) (Gibco; Cat.No. A1647801) according to the manufacturer’s instructions, and then subjected to single cell sequencing protocols. The differentiation of cortical neurons from NSCs was performed using doxycycline-inducible mNGN2. Neurons were then subjected to single cell sequencing protocols. Samples for single-cell 5’end RNA-seq were extracted using the Chromium Next GEM Single Cell 5’ Library and Gel Bead Kit v1.1 (PN: 1000165). Samples for snATAC-seq were extracted using the Chromium Next GEM Single cell ATAC Kit v1.1 (PN: 1000175), and sequenced via HiseqX Ten with the following conditions: R1: 150 cycles; R2: 150 cycles; Index1: 8 cycles; Index2: 8 cycles.

#### Definition of CREs from snATAC-seq data

The ATAC peak regions were defined by applying the Cell Ranger ATAC pipeline (10x Genomics) on the snATAC-seq sequencing files [Yip et al., in preparation]. The ATAC peaks were merged across all cell types, and represent the initial pool of genomic regions, *i.e.* CREs, further filtered and analyzed in this study. The ATAC signal across all the single cells was recounted using the fragment files by `CreateChromatinAssay` of the Signac package.

#### Transcriptional activity of CREs

From the single-cell 5’end RNA-seq, the output bam files generated by Cell Ranger [Yip et al., in preparation] were first provided to SCAFE (v1.0.1) [58] to identify genuine transcription start sites (TSSs). Briefly, genuine TSS clusters from each replicate were identified by `running scafe.workflow.sc.solo`. These clusters were then merged by running scafe.tool.cm.aggregate. Only the TSSs lying inside the genuine TSS clusters were used for quantifying CRE transcriptional activity, which accounted for 68.4% of all uniquely mapped reads. For each cell, the `scafe.tool.sc.count` function was used to count the amount of genuine TSSs signal within the CREs previously defined by ATAC-seq.

#### Metacell and pseudotime estimation

The quality control, clustering of cells into 391 metacells, and pseudotime inference is fully described in [Yip et al., in preparation]. Briefly SEACells [39] was used to group cells into 391 metacells according to the RNA and ATAC modalities. These 391 metacells, *i.e.* homogeneous cell clusters corresponding to granular cell types, were ordered along the differentiation pseudotime (Table S1).

#### Preparation of expression data and identification of transcriptionally variable CREs

To describe the variation of the CREs transcriptional activity along neuronal differentiation, the 5’-end RNA-seq counts pooled across metacells were prepared through the following steps:

1. Counts were normalized by metacell sizes
2. CREs with no expression signal in any of the metacells were discarded
3. The expression values were log-transformed after adding a small quantity equal to half of the minimum non-zero value of the data
4. CREs with a coefficient of variation lower than 0.5 were discarded
5. The expression values for each CRE were smoothed using GAMs and a spline component for the pseudotime
6. CREs with FDR-adjusted p-values greater than 1*e^−^*^5^ with respect to the spline component were discarded
7. The expression values for each CRE were scaled (z-score) in order to remove the differences in average expression intensity between CREs, and focus on their variation across metacells.

This resulted in the normalized smoothed expression profiles of 10912 transcriptionally variable CREs taken as input by STOIC (Table S2). To represent CREs in the reduced activity space, we rely on the tSNE algorithm, with a perplexity set to 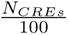, and other settings to their default value in the Rtsne package.

#### Extraction of CRE sequence features

The sequence of a CRE is defined by the center of the snATAC-seq peak extended by 500 bp upstream and 500 bp downstream. From these 1000 bp sequences, two types of sequence features were extracted.

- TFBM scores were computed from the non redundant PPM collection of JASPAR 2024 [38]. Motif occurrences were identified using FIMO (v5.5.1) from the MEME suite [59], with a p-value threshold of 1*e^−^*^4^, and by taking the maximum score on the two strands. When several occurrences of a motif were found in the same sequence, the motif with the highest score was kept. 744 PPMs had at least one occurrence in the CREs of this study. When no significant occurrence was found in a CRE, the score was set the the score of the minimum significant occurrence for this PPM minus 1. In the neuronal differentiation case study, we only kept the PPMs associated with a TF that is expressed in at least one metacell of the experiment, which is the case for 382 PPMs.
- K-mer frequencies were computed in each 1000 bp sequence for 1 ≤ *k* ≤ 3. The frequencies of a k-mer on the two strands are added, providing a total of 44 k-mer frequencies for each CRE.

### STOIC algorithm

STOIC proposes a clustering of the CREs based on their activity profiles, while maximizing the association between cluster membership and sequence features. STOIC takes as input two matrices:

- An activity matrix of dimensions *N_CREs_* ∗ *N_metacells_*. In non-single cell datasets, metacells are replaced by experimental conditions or time points.
- _•_ A matrix of sequence features of dimensions *N_CREs_* ∗ *N_DNAfeatures_*

The fully detailed STOIC algorithm can be found in Supplementary Methods A. Hereafter, we describe the main steps of the method. The clustering procedure of STOIC is composed of *N_start_* individual runs, defined by different random initializations, to improve stability.

#### STOIC run

The steps of an individual STOIC run are the following:

- **Step 1:** An initial centroid (*i.e.* a CRE) is randomly chosen in the activity profile space. All CREs with an activity profile correlated above a certain threshold (initially 0.8) with the profile of the centroid are included in the cluster. The centroid and the correlation threshold define the two parameters of the cluster.
- **Step 2:** A Random Forest (RF) [60] is fitted to predict the probability of each CRE to belong to the cluster using only sequence features as predictors. The AUC of the RF is estimated on Out-Of-Bag observations and stored. Then, the cluster parameters are optimized in order to increase the AUC. This involves slightly shifting, in turns, the cluster centroid and correlation threshold (see below): this modifies the set of CREs that belong to the cluster, and the AUC of the RF is re-estimated on this new cluster. If the AUC can be increased by such modifications, the RF is retrained on the updated cluster, and the whole process is repeated. Once the AUC can no longer be increased, and if the optimized cluster has an AUC higher than a user-defined AUC threshold, it is added to the list of optimized clusters. The total CRE coverage, *i.e.* the fraction of CREs covered by a cluster of the list is computed. Then, a new centroid is randomly picked far from the initial and final centroids of the optimized clusters, and step 2 is repeated.
- **Step 3:** Once the CRE coverage stops increasing, the run is over and the optimized clusters and their supervised models are returned.

##### Optimization of the correlation threshold

A series of 8 candidate correlation thresholds spread between the current correlation threshold ±20% are tested. The first candidate correlation that improves the AUC after re-training the RF is selected to replace the former threshold.

##### Optimization of the cluster centroid

All CREs belonging to the current cluster are tested as a new centroid (to speed up the process, if the current cluster contains more than 500 CREs, only the most correlated 500 CREs are tested). The first candidate centroid that improves the AUC after re-training the RF is selected to replace the former centroid. The number of RF re-fits is limited to 10 for the sake of speed.

#### Clusters pruning and CREs assignation

Once all *N_start_*runs have been performed, the clusters from the different runs are aggregated and pruned. Indeed, several clusters may converge to the same area of the activity space and exhibit similar sequence features. In order to get a set of final, non-redundant clusters, the cluster list is pruned based on cluster overlap (*i.e.*the amount of CREs shared by two different clusters, as measured by the Jaccard index), the correlation of activity profiles, and a ”swap score”. The swap score of two clusters measures the similarity of the RFs associated with these clusters. It is defined by min(swap_12_, swap_21_ ) with 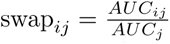, *AUC_ij_* the AUC of model *i* for predicting the classes formed by cluster *j*, and *AUC_j_* the AUC of model *j* for predicting cluster *j*. When two clusters are deemed redundant based on these three metrics, only the one with the best AUC is kept. When the pruning has been done, the CREs that belong to two clusters or more are assigned to a single one. Indeed, even after pruning, it can happen that two clusters overlap. In this case, the shared CREs are assigned to the cluster where they rank higher in terms of posterior probability, *i.e.* they become members of the most likely cluster given their sequence features.

#### Computation of feature importance

Finally, the importance of each sequence feature is computed based on the supervised model trained on each cluster. Two options are available in STOIC to compute feature importance:

- **Permutation importance**, measured by the drop in prediction performance caused by shuffling a feature *X_j_* on Out-Of-Bag observations as originally proposed by L. Breiman when introducing Random Forests [37].
- **Conditional permutation importance**, an extension of permutation importance with the objective of reducing bias caused by correlation between features [61]. In this approach, the shuffling of Out-Of-Bag observations is performed within partitions of the values of the predictors in each tree that are correlated with *X_j_* over a certain threshold (set to 50%) and under a certain depth level in the trees (set to 8).

On the neuronal differentiation dataset, STOIC was run with 100 different initializations, an AUC cut-off of 0.7 and 500 trees.

### Comparison to alternative clustering approaches

To assess the interest of our guided clustering approach, two alternative strategies based on co-expression clustering were investigated.

#### K-means

K-means clusters were computed for different numbers of clusters (4 ≤ *k* ≤ 16). As the k-means is known to be sensitive to initialization randomness, for each number of clusters, 100 kmeans were run with different initialization, and the best clustering in terms of the within cluster sum of square (*i.e.* the statistic optimized by the kmeans algorithm) was kept.

#### hdWGCNA

HdWGCNA [33] is an adaptation of the original WGCNA algorithm [62] to single-cell and spatial omics. We used as input our raw CREs per metacell expression assay, prepared with the `SetupForWGCNA()` and `SetDatExpr()` functions with default parameters and by defining a custom set of genes, *i.e.* the CREs of our study. The `ConstructNetwork()` function was then used to build the co-expression modules with default parameters, and the soft power parameter automatically estimated by a call to `TestSoftPowers()`.

#### Coverage correction

In STOIC and the hdWGCNA algorithm (but not in kmeans), all CREs are not assigned to a specific cluster, and some CRE remain unclassified. In order to obtain comparable clustering for the three approaches, a coverage correction was applied. Given the STOIC clustering as reference because it is the one with the highest number of unclassified CREs, the procedure involves to iteratively remove from the kmeans and the hdWGCNA clustering the CRE that is the furthest away from its centroid, and to repeat the operation until the same coverage as in STOIC is reached.

### *De-novo* motif discovery with STREME

For each cluster, ^2^ of the positive and negative CRE examples were used for *de-novo* motif learning with STREME (v5.5.1, default options). Test CREs consisting of the remaining ^1^ of CREs were then scanned with the *de-novo* discovered motifs, using FIMO (v5.5.1) with a p-value threshold of 1*e^−^*^4^, taking the maximum score on the two strands, and keeping the motif with the highest score per CRE. The AUCs of RFs leveraging k-mers frequencies, JASPAR motifs, and/or the *de-novo* motifs scores were then computed and compared on the test CREs only.

### DNA sequence features heatmaps

Features heatmaps show the sequence feature scores of the most important features (columns) in the CREs of these clusters (rows). The restricted list of most important features was obtained via the ”elbow” rule from the feature importance curve. This elbow method is implemented in the find curve elbow function from the pathviewr R package. Important features were restricted beforehand to the features with higher scores in the cluster than in other CREs. The scores of each DNA feature were scaled to a z-score across the 10912 CREs, so that color scales between columns are comparable. Red cells represent scores higher than average, and blue scores below than average. Heatmaps were represented using the ComplexHeatmap Bioconductor package.

### Simulations

We tested STOIC on datasets designed to contain co-expression clusters associated with specific, known sequence features. Simulating such a ground truth allows to assess STOIC’s ability to retrieve known associations, and its sensitivity to varying intensities of sequence features enrichment. The true neuronal differentiation dataset served as a basis for generating artificial datasets. The expression data of the CREs in the different metacells was left untouched. Co-expression clusters were formed using k-means (*k* = 10). On the other hand, we shuffled independently all the sequence features (column-wise permutations) to break the link between CREs and any of their sequence characteristics. Then, we artificially enriched one or several sequence features in the CREs of some of the k-means co-expression clusters. Two clusters received an enrichment in a single TFBM (respectively NEUROD1 and RFX3), two clusters received an enrichment in a single k-mer (respectively AGA and GC), one cluster received an enrichment in both the ACT k-mer and the TFBM of MAF, and one cluster received an enrichment in the product of two TFBMs scores (TEAD1 and GLI3) which simulates an interaction effect. 4 clusters did not receive any enrichment. The intensity with which enrichments were performed is a parameter of the simulations (*e*), with possible values between 0 and 1. To enrich a feature in a given co-expression cluster of size *S*, we assign the *e* ∗ *S* feature scores with the highest values to *e* ∗ *S* CREs sample from this cluster. The remaining (1 − *e*) ∗ *S* CREs from this cluster and all other CREs share the *N* − *e* ∗ *S* lowest values for this feature. At *e* = 0 there is no enrichment. At *e* = 1, the enrichment is maximal as all CREs of the cluster have a feature value higher than all CREs outside the cluster.

We ran STOIC with 500 trees, an AUC threshold of 0.6, and 10 starts. To quantify the performance of STOIC on these simulations, the (adjusted) Rand Index between STOIC’s inferred clusters and the ground truth clusters was computed. The important sequence features of STOIC’s clusters were also be compared to artificially enriched one(s) in the known clusters.

All code and notebooks related to these simulations can be found at https://gite.lirmm.fr/ml4reggen/stoicsimulations.

### CUT&Tag dataset for CREs annotation

The methods of CUT&Tag chromatin state data, generated in iPSCs, NSCs, and neurons were described in the FANTOM6 publication (Yip et al. 2024). The extraction protocol and bio-informatic pre-processing are detailed in Supplementary Methods B. Chromatin states of CREs were defined in each of the 3 cell-types by ChromHMM [63] using the CUT&Tag epigenetic marks and scATAC-seq data based on 200-bp genomic bins. This resulted in 16 clusters that were manually annotated according to their chromatin features and proximity to TSSs. The derived chromatin states, *i.e.*promoter (promoter and promoter flanked), bivalent promoter, active enhancer, repressed enhancer, primed enhancer, and CTCF-alone were then intersected with the CRE regions in each cell-type. Finally, by combining chromatin states in the 3 cell types, the 10912 CREs of this study were broadly classified as “promoter-like”, ”enhancer-like”, “CTCF-alone” or “unclassed”. Based on the extremely low numbers of “CTCF-alone” and “unclassed” (resp. 15 and 65 CREs), only the “promoter-like” and ”enhancer-like” signatures were discussed in the results.

### ChIP-Seq dataset

ChIP-Seq datasets of TEAD4 in iPSCs and ONECUT2 in neurons were generated according to the extraction, sequencing and bio-informatic protocols in Supplementary Methods B. We considered only the ChIP-Seq peaks containing the motif of the corresponding TF. We then assessed the enrichment in binding events in the different CRE clusters based on these two experiments. For each cluster, the number of CREs overlapping a peak in at least one of the ChIP-Seq replicate was computed, and enrichment for binding events within the cluster was then tested by a one-sided hyper-geometric test and all CREs as background. We restricted this analysis to the CREs correctly assigned to their cluster by STOIC’s supervised models, *i.e.*their sequence features correctly predict them as members of the cluster.

### CRE to genes annotations

A CRE was associated to a gene model if the 5’-end of the gene’s transcript model is initiated within the CRE region, without extension.

### Repeat elements analysis

Repeat elements were taken from the RepeatMasker annotation (https://genome.ucsc. edu/cgi-bin/hgTables, group: ’Repeats’, track: ’RepeatMasker’), and intersected with the CREs of this study. We restricted this analysis to the CREs correctly assigned to their cluster by STOIC’s supervised models, *i.e.*their sequence features predict them as members of the cluster. The types of repeat elements considered were repeat Classes (e.g SINE, LINE, LTR, . . . ) or the finer-grained repeat Names (e.g HERV-int, LTR7, . . . ). For each cluster and each type of repeat element, a fisher’s exact test was performed. The enrichment p-values were adjusted by the Benjamini-Hochberg false discovery rate correction.

### Interactome datasets

#### Hi-C dataset

Hi-C in iPSCs, NSCs and neurons was performed as described in [64], with details provided in Supplementary Methods B. The pre-processed Hi-C data was then transformed into valid genomic interaction pairs at 5 kb resolution. The statistical significance of these interactions was calculated using the Bioconductor package GOTHiC (version 1.40.0). Only intra-chromosomal (cis) interactions supported by at least five read pairs and a q-value (FDR) ≤ 0.01 were deemed significant for downstream analysis.

The Hi-C interactions were then intersected with the CREs of this study and interactions involving the same CRE were removed. The intra-cluster interaction rate of a cluster *C*_1_ is computed as

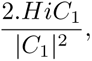

with *HiC*_1_ the number of Hi-C interactions between CREs of *C*_1_. The inter-cluster interaction rate between different clusters *C*_1_ and *C*_2_ is computed as

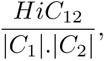

with *HiC*_12_ the number of Hi-C interactions between CREs of *C*_1_ and *C*_2_.

#### RADICL-Seq dataset

The protcol used to generate the RNA-DNA interactions data with RADICL-Seq in iPSCs, NSCs and Neurons is available in [Pracana et al., in preparation]. This dataset was further characterised in [Pracana et al., in preparation, Lambolez et al., in preparation]. The RADICL-Seq interactions were intersected with the CREs of this study to obtain a CRE-CRE interaction table. Interactions involving the same CRE from the RNA and DNA side were removed as they represent transcription events rather than other forms of regulatory mechanisms. The intra-cluster interaction rate of a cluster *C*_1_ is computed as

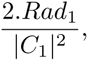

with *Rad*_1_ the number of interactions between RNAs and DNAs from CREs of *C*_1_. There are two different inter-cluster interaction rates between clusters *C*_1_ and *C*_2_ :

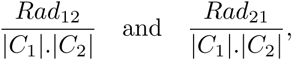

with *Rad*_12_ the number of interactions between the RNAs of CREs from *C*_1_ the DNAs of CREs from *C*_2_ and *Rad*_21_ the number of interactions between the RNAs of CREs from *C*_2_ the DNAs of CREs from *C*_1_.

### List of intellectual disability genes

We retrieved the list of genes involved in neurodevelopmental disorders from the sysNDD application at https://sysndd.dbmr.unibe.ch/.

### Application of STOIC to bulk FANTOM5 CAGE kinetic experiments

The expression data of the kinetic experiments described in [13] was retrieved from https://fantom.gsc.riken.jp/5/datafiles/latest/extra/CAGE_peaks/hg19.cage_peak_phase1and2combined_counts_ann.osc.txt.gz (for promoters) and https://fantom.gsc.riken.jp/5/datafiles/latest/extra/Enhancers/human_permissive_enhancers_phase_1_and_2_expression_count_matrix.txt.gz (for enhancers). Expression data was pre-processed with the following steps:

1. Counts were normalized by sample size
2. The profile of each CRE and replicate was smoothed using a rolling mean of window 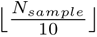
3. The profile of each CRE and replicate was scaled with a z-score

The sets of transcriptionally variable CREs for the different time courses are the ones identified in [13], retrieved from Supp. Table 3. This table also provided the CRE annotation (enhancer, TF promoter or gene promoter). The sequence features were extracted in the same manner as in the neuronal differentiation dataset, *i.e.* non-overlapping 1000 bp sequences were formed for each CRE. In this case, CRE sequences were centered around the CAGE peak center instead of the ATAC-seq peak center, and the PWMs were restricted to the PPMs of the TFs that are transcriptionally variable in the corresponding time course. STOIC was run with 20 different initializations for each time series, an AUC cut-off at 0.65, and 500 trees. All R scripts used to prepare data, run STOIC and generate results are available in the repository https://gite.lirmm.fr/ml4reggen/stoic f5 kinetics.

## Supplementary information

This article is accompanied by:

- One Supplementary Tables file,
- One Supplementary Figures file,
- One Supplementary Methods file.

Supplementary Methods A contains the detailed STOIC algorithm. Supplementary Methods B contains detailed experimental protocols for data generation and upstream dry analysis.

The code, data and notebooks needed to run STOIC and reproduce the analyzes of the paper can be found in the following repositories:

- STOIC R package: https://gite.lirmm.fr/ml4reggen/stoic
- The application of STOIC to neuron differentiation: https://gite.lirmm.fr/ ml4reggen/stoic neuron diff
- Simulations: https://gite.lirmm.fr/ml4reggen/stoic simulations
- The application of STOIC to bulk CAGE time courses: https://gite.lirmm.fr/ ml4reggen/stoic f5 kinetics

## Acknowledgements

We are grateful to the ML4REGGEN team for helpful discussions, as well as Shaun Mahony and Marcel Schulz.

## Declarations

### Ethics approval and consent to participate

Not applicable.

### Consent for publication

All authors have read the article and give their consent for publication.

### Author contribution

**Oćeane Cassan**: Conceptualization, Investigation, Computational analysis, Project supervision, Writing (original draft), Writing (review & editing)

**Julien Raynal**: Computational analysis, Investigation, Writing (original draft)

**Christophe Vroland**: Computational analysis, Investigation

**Kayoko Yasuzawa**: Experimental analysis

**Tsukasa Kouno**: Experimental analysis

**Jen-Chien Chang**: Computational analysis

**Chung-Chau Hon**: Conceptualization, Investigation, Computational analysis

**Jay W. Shin**: Project supervision, Funding acquisition

**Masaki Kato**: Project administration

**Hazuki Takahashi**: Project administration

**Takeya Kasukawa**: Database management

**Nobusada Tomoe**: Database management

**Vincenzo Lagani**: Investigation, Writing (review & editing)

**Robert Lehman**: Investigation

**Kévin Yauy**: Investigation, Computational analysis

**Piero Carninci**: Project administration

**Chi Wai Yip**: Conceptualization, Investigation, Computational analysis, Project supervision, Project administration, Funding acquisition, Writing (review & editing)

**Laurent Bréhélin**: Conceptualization, Investigation, Computational analysis, Project supervision, Project administration, Funding acquisition, Writing (original draft), Writing (review & editing)

**Charles-Henri Lecellier**: Conceptualization, Investigation, Computational analysis, Project supervision, Project administration, Funding acquisition, Writing (original draft), Writing (review & editing)

