## Supplementary Tables Caption for "A genome-wide, machine learning-guided exploration of the *cis*-regulatory code involved in neuronal differentiation"

**Table S1**: **Pseudotime information.** The inferred pseudotime used to order metacells in the neuronal differentiation process.

**Table S2**: **CRE activity matrix.** The normalized, scaled, smoothed expression (scRNA-Seq) of transcriptionally variable CREs used as input for STOIC.

**Table S3**: **BED file of CREs.** The genomic coordinates (hg38) of length 1000 bp used to extract CREs sequence features.

**Table S4**: **CREs sequence features matrix.** The 382 TFBM scores and 44 k-mers frequencies of the CREs used as input for STOIC.

**Table S5**: **CREs clusters identified by STOIC.** For each CRE, the cluster from STOIC is given along with the "well predicted" information (whether or not the CRE is correctly predicted as a member of the cluster by the supervised model of STOIC).

**Table S6**: **Important sequence features in STOIC clusters**. For each cluster, the 20 most important sequence features (TFBMs or k-mers) extracted from RF models, ranked by conditional permutation importance.

**Table S7**: **CREs annotation.** For each CRE, the epigenetic signature (e.g. enhancer, promoter, . . . ) as predicted by chromHMM based on CUT&Tag data.

**Table S8**: **TF expression profiles.** The normalized, scaled, smoothed expression (scRNA-seq) in the different metacells for the TFs corresponding to the 382 TFBMs included as sequence features in this study.

**Table S9**: **CHIP-Seq dataset for TEAD4.** Intersection between TEAD4 CHIP-seq peaks and CREs, with information about motif occurrence within peaks for TEAD1/2/3/4.

**Table S10**: **CHIP-Seq dataset for ONECUT2.** Intersection between ONECUT2 CHIP-Seq peaks and CREs, with information about motif occurrence within peaks for ONECUT1/2/3.

**Table S11**: **Significant cluster enrichments in repeat classes from RepeatMasker.** Each significant enrichment for a given cluster and a given repeat Class after FDR correction is reported, along with the number of CREs with the repeat in the cluster.

**Table S12**: **Significant cluster enrichments in repeat names from RepeatMasker.** Each significant enrichment for a given cluster and a given repeat Name after FDR correction is reported, along with the number of CREs with the repeat in the cluster.
