## Supplementary Figures for "A genome-wide, machine learning-guided exploration of the *cis*-regulatory code involved in neuronal differentiation"

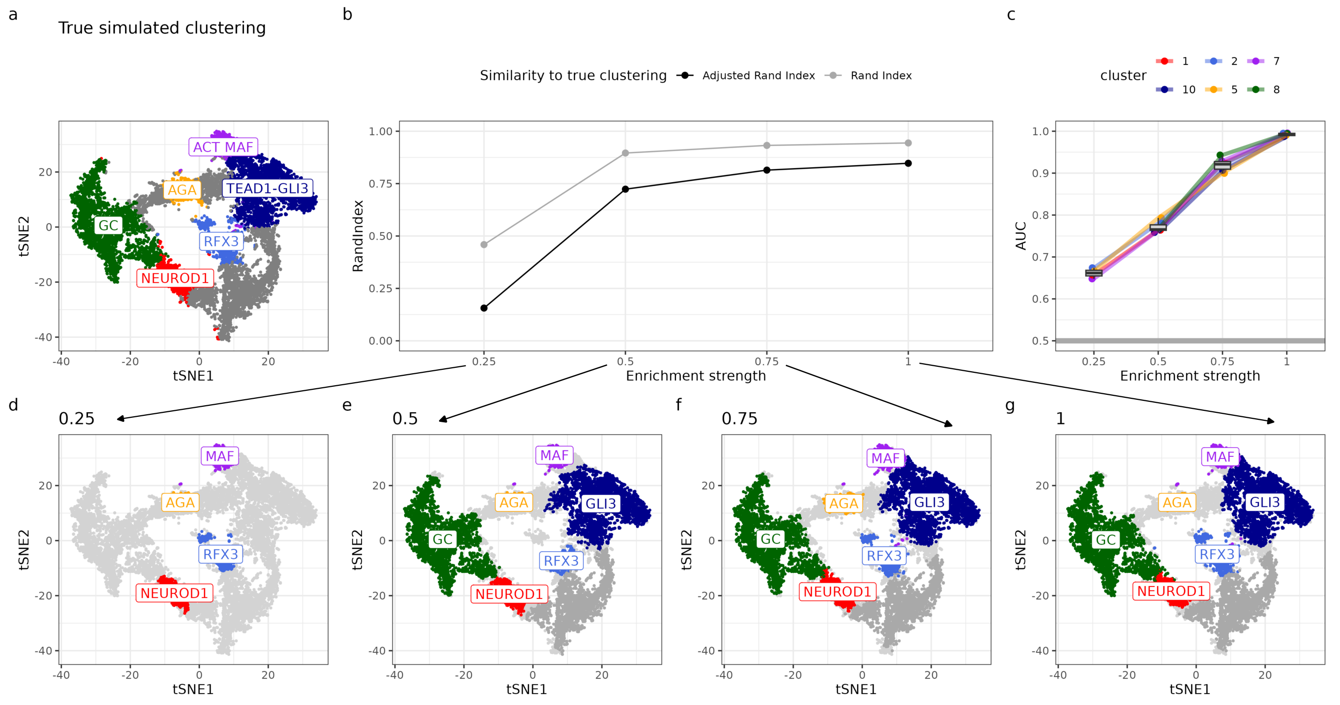


**Fig. S1**: **STOIC retrieves a simulated gold standard of co-activity clusters and enriched DNA features.** **a.** Ground truth of co-activity clusters where sequence features were artificially enriched, represented in the reduced activity space via a tSNE. **b.** Similarity to the ground truth clusters (Rand Index and Adjusted Rand Index) as a function of DNA features enrichment. **c.** AUC of inferred clusters as a function of DNA features enrichment. **d-g** Clusters retrieved by STOIC and the associated most important DNA features. Each panel represents a different enrichment strength for DNA features.

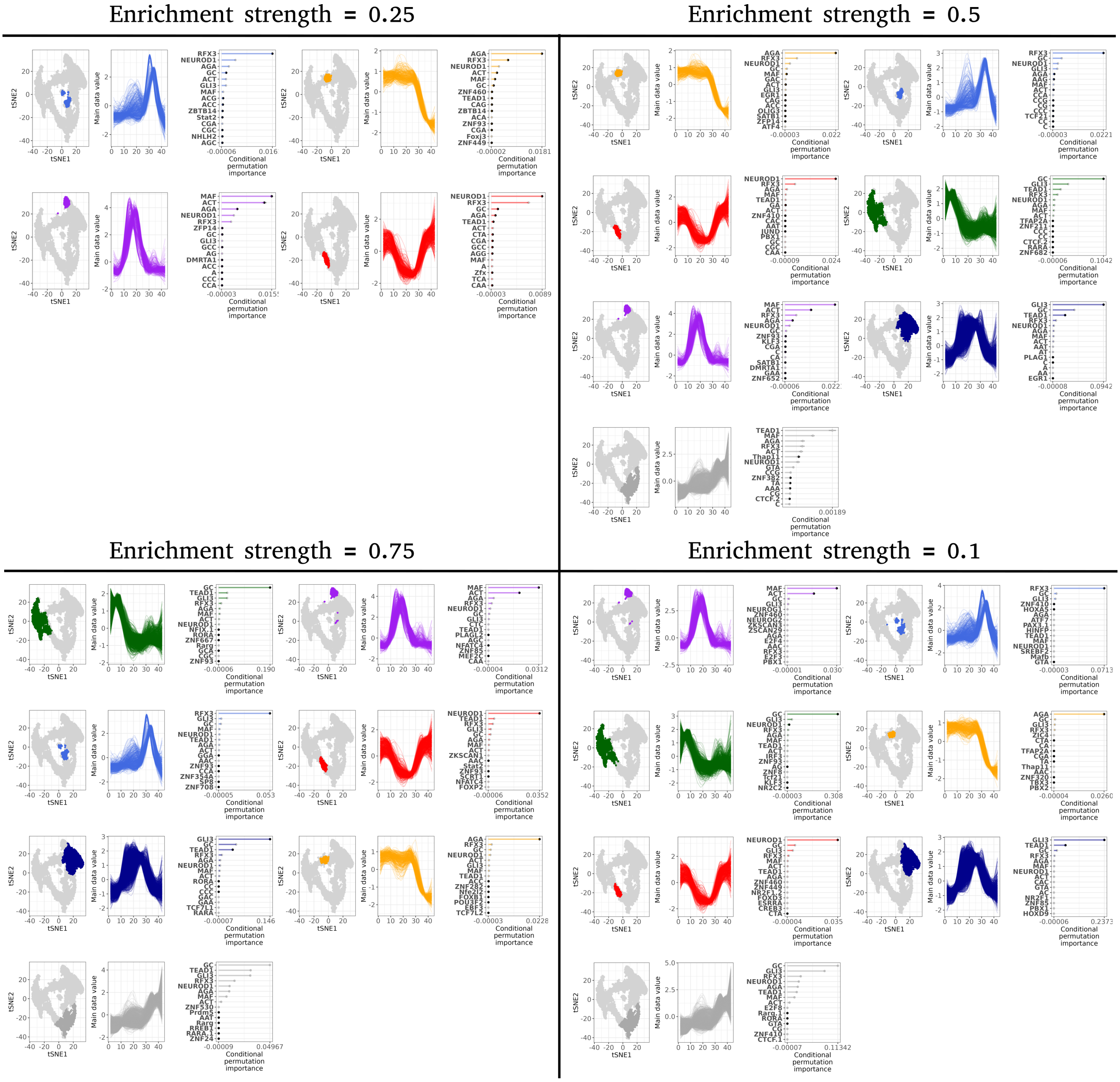


**Fig. S2**: **Detailed simulation results illustrate STOIC clusters and sequence features on a simulated ground truth.** For each enrichment strength and each inferred cluster: the cluster in the reduced activity space, the cluster activity profile, and the most important sequence features.

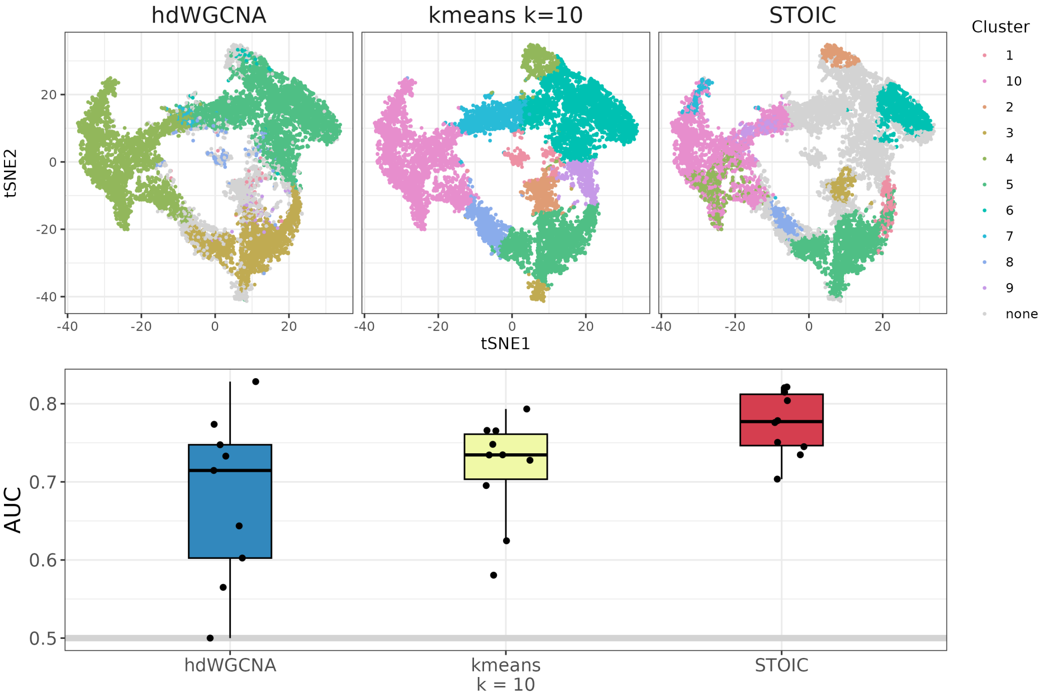


**Fig. S3**: **STOIC recovers more predictive sequence features than alternative co-activity clustering approaches without coverage adjustment.** On the first row, the co-activity clusters of the different approaches: hdWGCNA, k-means with $k = 10$, and STOIC represented in the reduced activity space via a tSNE. On the second row, the AUC achieved when predicting cluster membership using CREs sequence features.

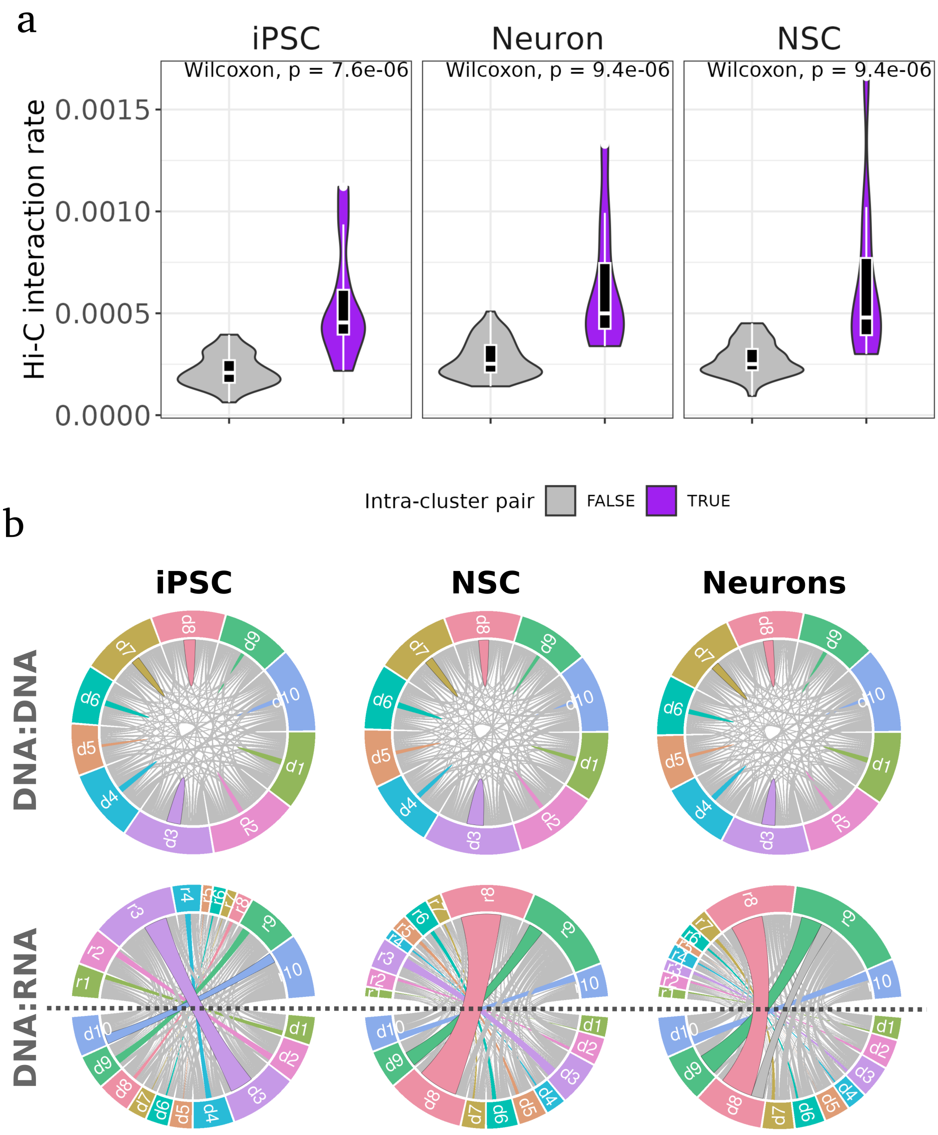


**Fig. S4**: **DNA:DNA and DNA:RNA interactions rates between STOIC clusters. a.** DNA:DNA interaction rates derived from Hi-C data in iPSCs, NSCs, and neurons between pairs of STOIC clusters. Purple violins denote CRE interaction rates within the same cluster, whereas grey violins denote CRE interaction rates between two different clusters. **b.** Interaction rates in the Hi-C (first row) and RADICL-Seq (second row) datasets. Self-interacting CREs were removed. The clusters represent the different sectors, and the width of links represents the rate of interactions between CREs of the two related clusters. Only intra-cluster interaction rates are colored. "d": DNA side of an interaction, "r": RNA side of an interaction.

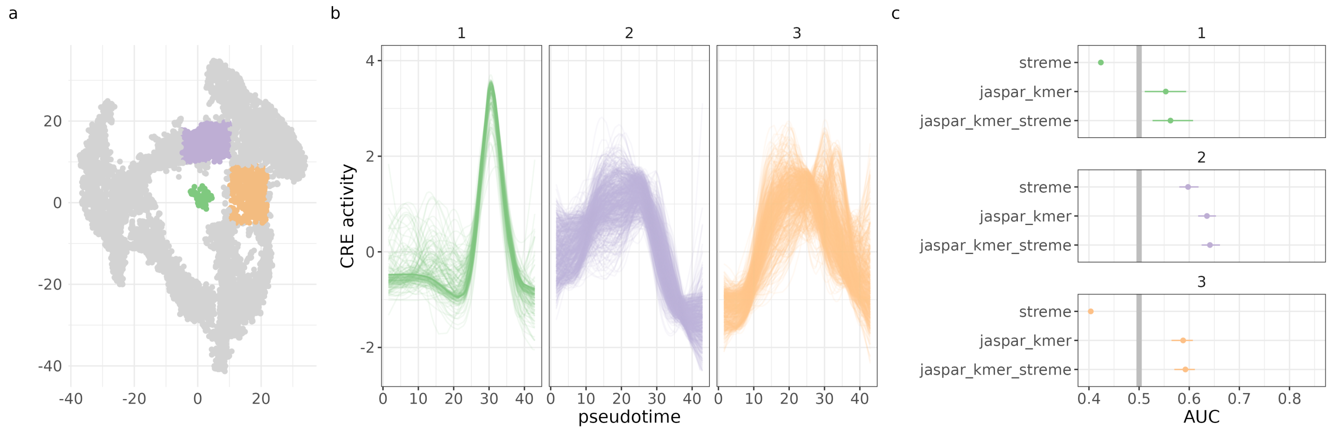


**Fig. S5**: **STOIC leaves out certain areas of the activity space because of low AUCs. a.** Areas in which STOIC found no clusters, represented in the reduced tSNE space. b. Activity profiles of the areas that were not retrieved by STOIC. c. AUC of different sequence-based RF models to predict the areas that were not retrieved by STOIC. "streme": *de-novo* STREME motifs alone, "jaspar_kmer": TFBMs and k-mers frequencies, "jaspar_kmer_streme": TFBMs, k-mers frequencies and *de-novo* STEME motifs.

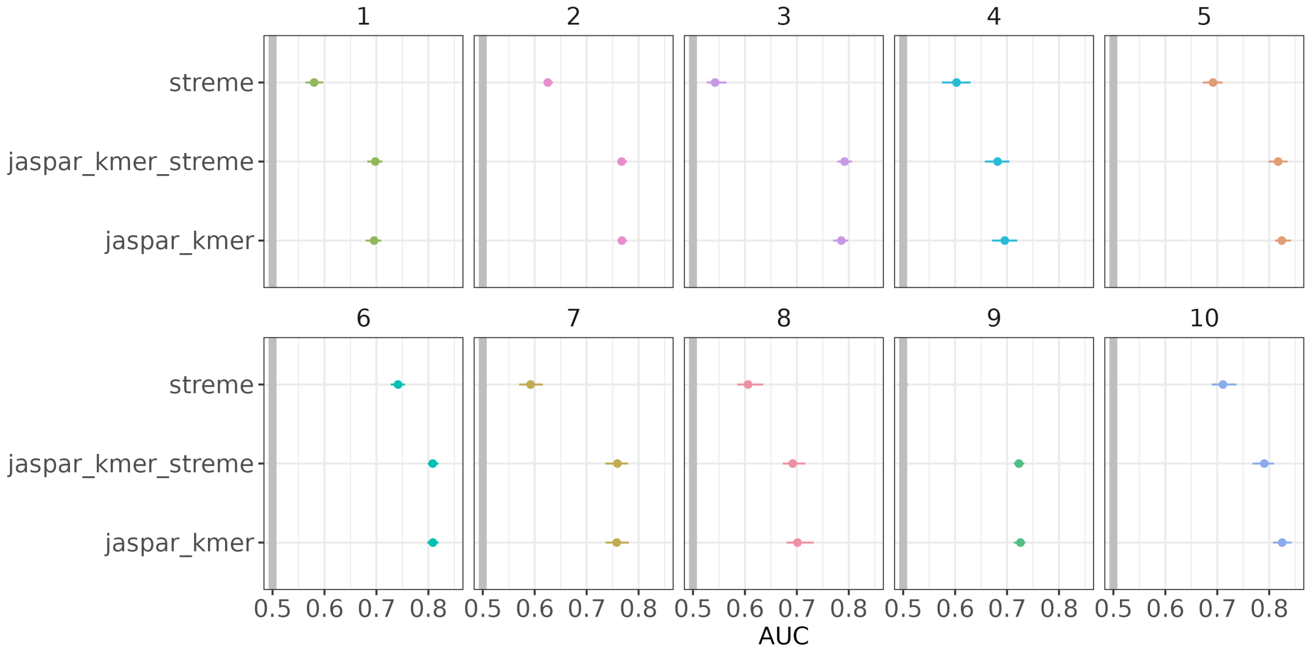


**Fig. S6**: ***De-novo*** **STREME motifs do not improve the AUCs of STOIC clusters. a.** AUC of different sequence-based RF models to predict STOIC clusters. "streme": *de-novo* STREME motifs alone, "jaspar_kmer": TFBMs and k-mers frequencies, "jaspar_kmer_streme": TFBMs, k-mers frequencies and *de-novo* STEME motifs.

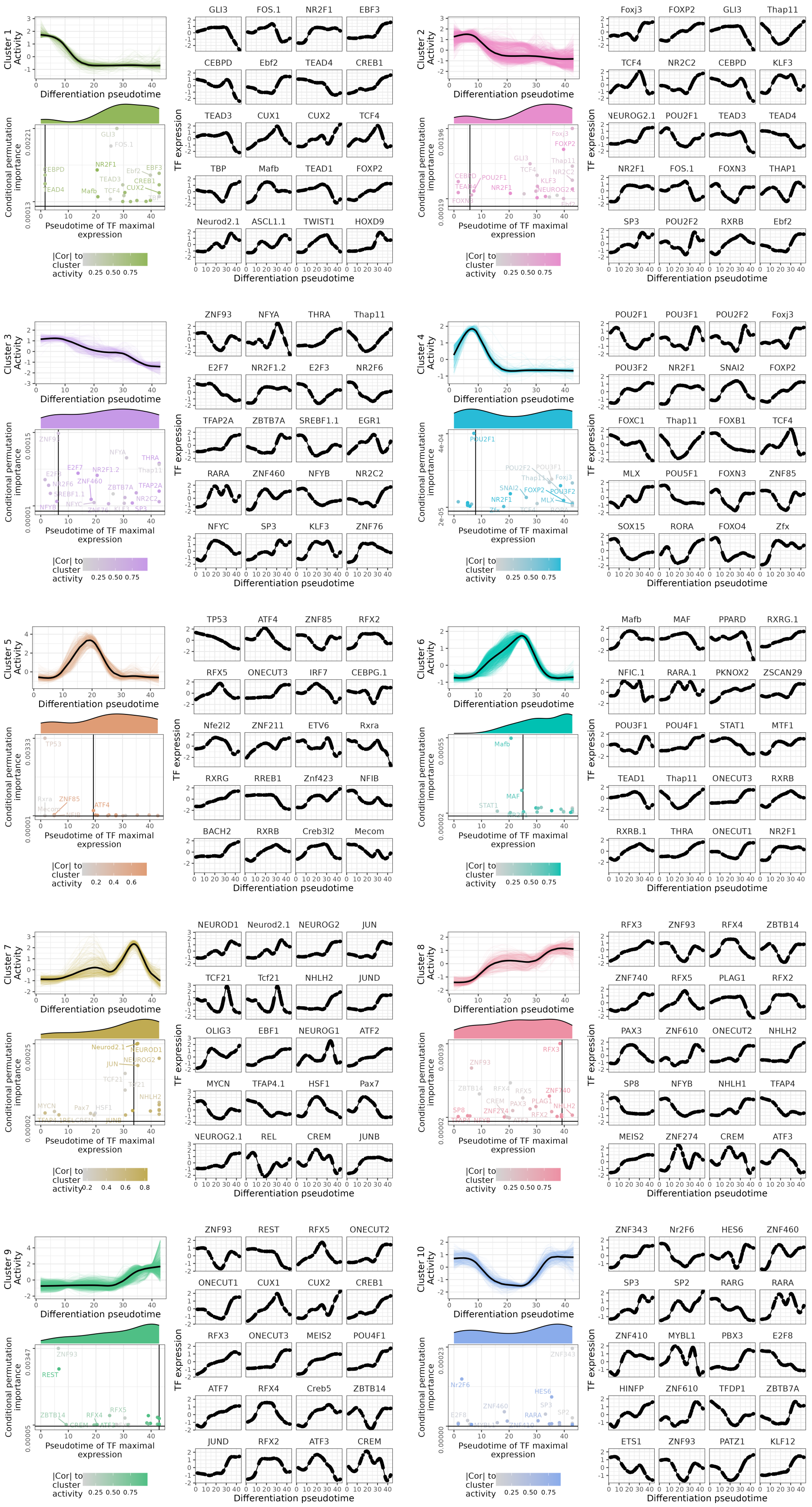


**Fig. S7**: **Expression dynamics of the 20 most important TFs in each cluster.** Each panel shows a cluster activity profile, and the conditional permutation importance as a function of the TF pseudotime of maximal expression, for the 20 most important TFs in the cluster. Important TFs are restricted to the TFs with higher scores in the cluster (i.e. positively associated to the CREs of the cluster). The vertical line represents the pseudotime of maximal activity of the co-activity cluster. The marginal distributions of the TFs pseudotimes of maximal expression are shown above the scatter plot. TFs are colored depending on the absolute correlation of their expression profile to the activity profile of the cluster. For each cluster, we also report the detailed expression profile of the 20 most important TFs. TF expression is normalized, smoothed and scaled to a z-score.

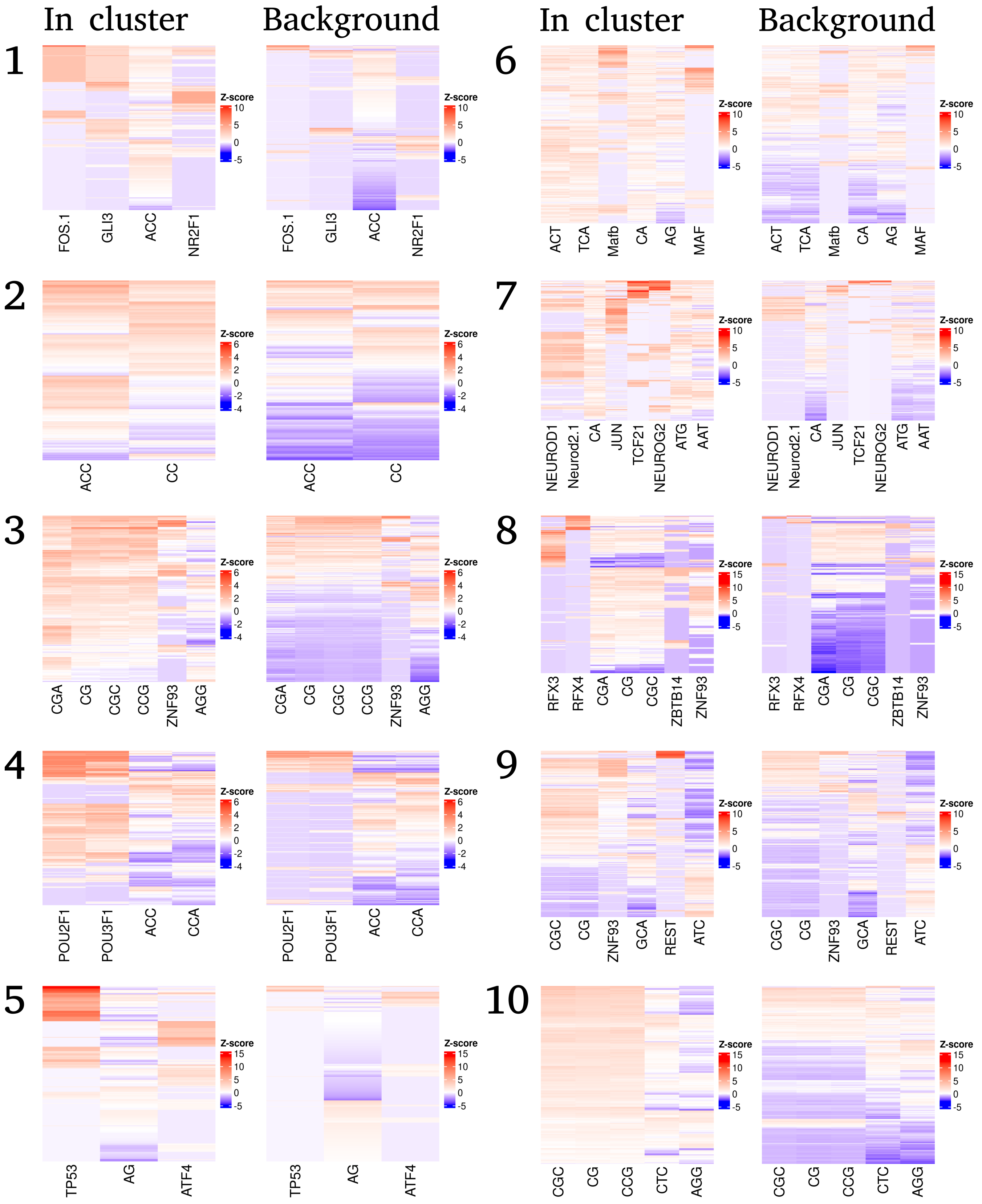


**Fig. S8**: **Distribution of important DNA features scores in STOIC clusters.** Heatmap of sequence feature scores of the most important features (columns) in the CREs (rows). The heatmap on the left shows the CREs of the cluster, and the heatmap on the right shows the other CREs. The restricted list of most important features was obtained via the "elbow" rule from the ranked features importance curve. Important features were restricted beforehand to the features with higher scores in the cluster than in other CREs. The scores of each DNA feature were scaled to a z-score across the 10912 CREs. Red cells represent scores higher than average, and blue scores below than average.

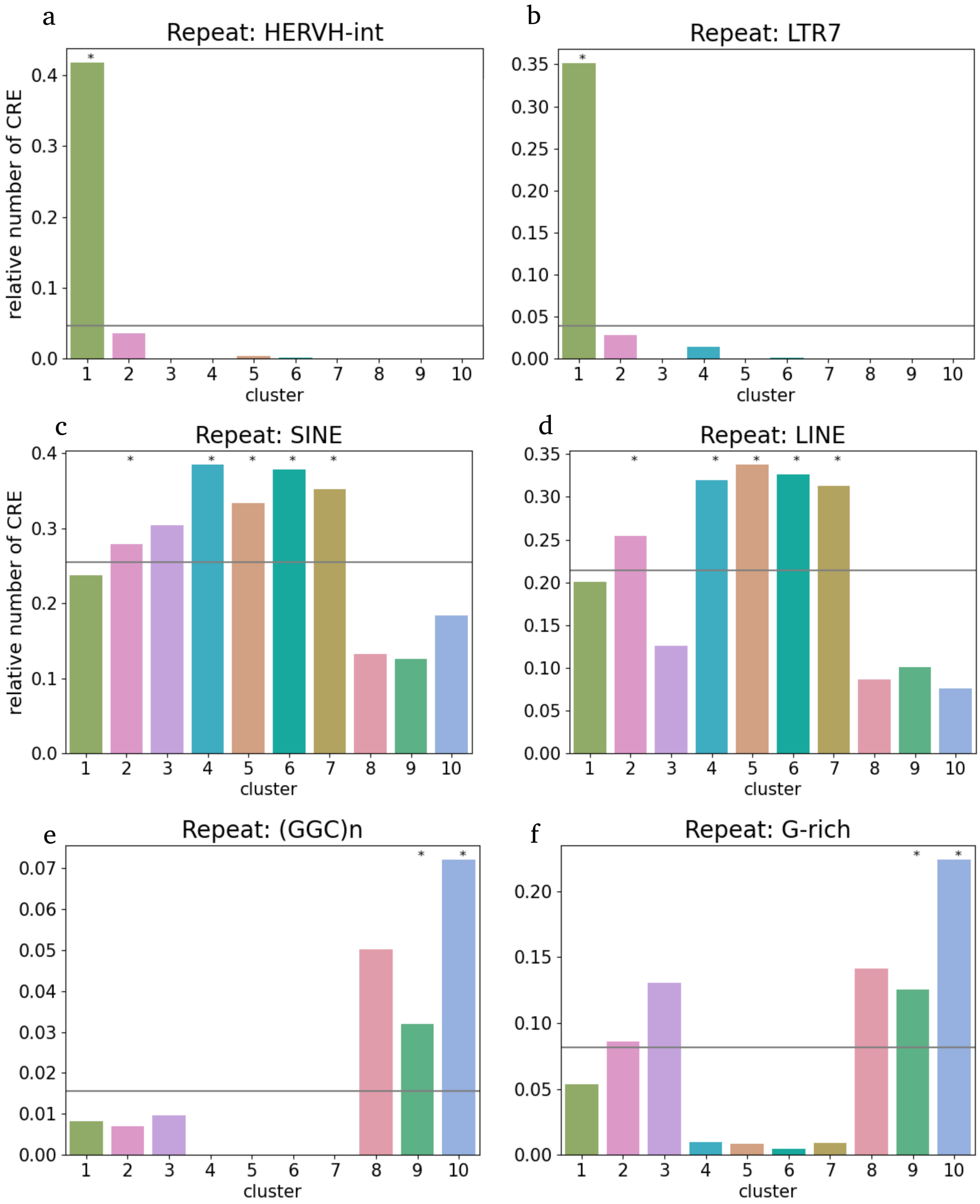


**Fig. S9**: **The CREs of different clusters are enriched in distinct repeat elements. a-f.** Fraction of CREs in each cluster overlapping different types of repeats from the RepeatMasker annotation. Stars denote a significant enrichment ($P < 0.05$) of repeats based on a Fisher's exact test after FDR correction. The grey bar represents the baseline proportion.

| GENE | GC-REPEAT EXPANSION DISEASE | SYSNDD DISEASE | CLUSTER | CRE |
|---|---|---|---|---|
| PPP2R2B | SCA12 | - | 8 | chr5-146877621-146878229 |
| ATXN8OS | SCA8 | | 9 | chr13-70107439-70108619 |
| AFF2 | FRAXE | Intellectual developmental disorder, X-linked 109 | 9 | chrX-148500363-148501701 |
| TCF4 | FECD3 | Pitt-Hopkins syndrome | none | chr18-55321613-55322814;chr1[illegible] 55401514-55402501 |
| DMPK | DM1 | Myotonic dystrophy 1 | none | chr19-45781982-45782867 |
| AFF3 | FRA2A | KINSSHIP syndrome, intellectual disability | none | chr2-100192059-100192727 |

**Fig. S10**: **Intersection of the 28 genes associated with neuropathologies via GC-rich repeat expansions and STOIC clusters.** Gene symbols are shown along with the GC-repeat expansion disease, the sysNDD disease if available, the cluster in which the gene transcript is initiated, and the CRE in which the gene is initiated.

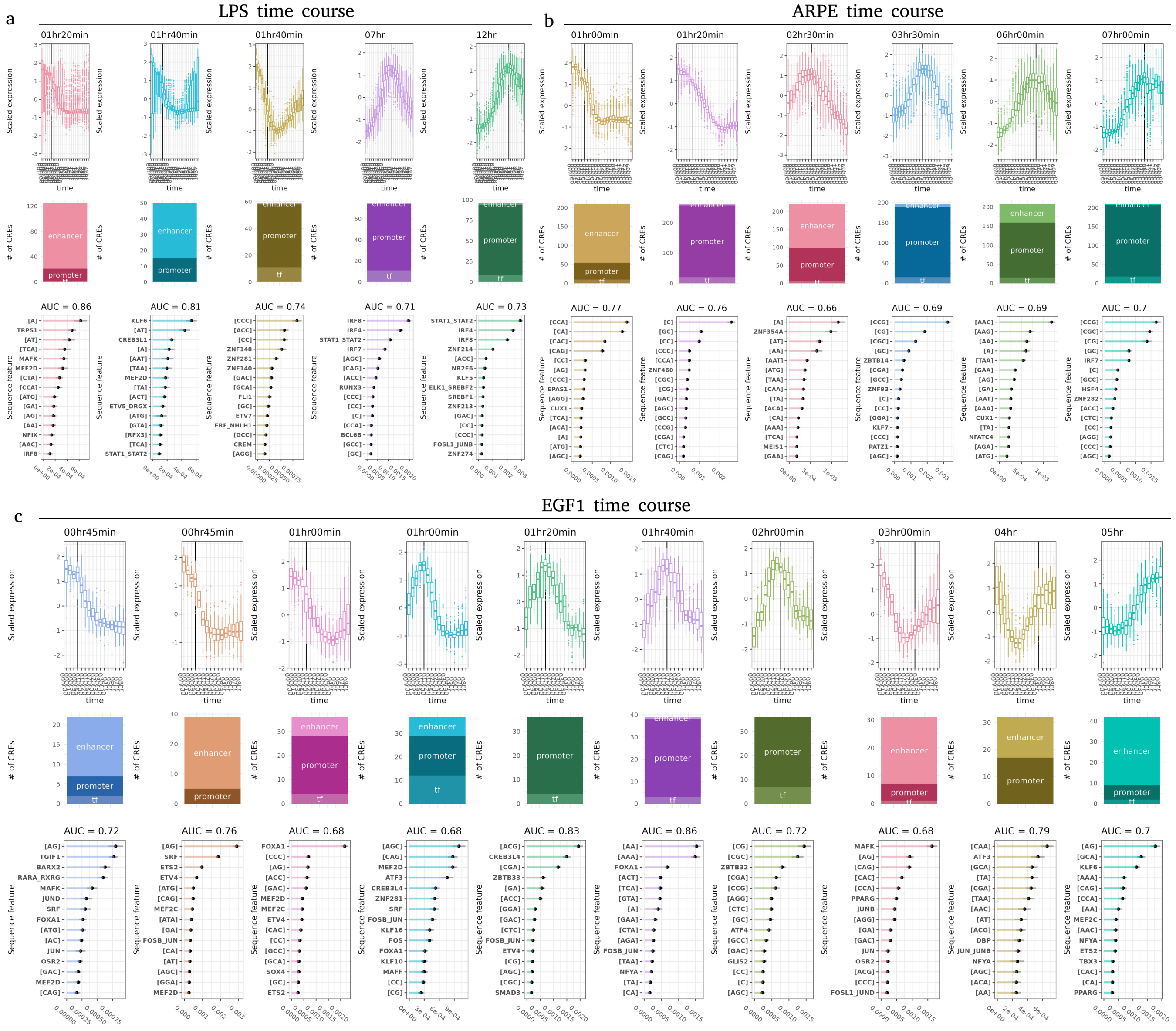


**Fig. S11: Detailed STOIC results on the LPS, ARPE and EGF1 bulk CAGE time courses. a-c.** In each kinetic experiment, detailed results include the normalized activity profiles of each cluster (1st row), the composition of the cluster (2nd row), and the important sequence features and AUC in each cluster (3rd row). The vertical line in the activity profiles corresponds to the profile center of mass, i.e the time point where half of the CRE activity has changed. Clusters are ordered by center of mass. CRE annotation: "enhancer", "tf" (for a TF promoter), "promoter" (for the promoter of a non-TF gene).
