## Supplementary Methods for "A genome-wide, machine learning-guided exploration of the *cis*-regulatory code involved in neuronal differentiation"

### A. Detailed STOIC algorithm

STOIC proposes a clustering of the CREs based on their activity profiles, while maximizing the association between cluster membership and sequence features. STOIC takes as input two matrices:

- An activity matrix of dimensions $N_{CREs} * N_{metacells}$. In non-single cell datasets, metacells are replaced by experimental conditions or time points.
- A matrix of sequence features of dimensions $N_{CREs} * N_{DNAfeatures}$

The clustering procedure of STOIC is composed of $N_{start}$ runs to improve the stability and robustness of the results. Each run is defined by a different random initialization and returns a collection of clusters of co-active CREs and their supervised model learned on sequence features (see 0.1). Once all $N_{start}$ runs have been performed, the clusters from the different runs are aggregated and pruned (see 0.5), and CREs are dispatched between all clusters (see 0.6). Finally, sequence feature importance (see 0.7) is computed using the supervised model train on each cluster. STOIC returns the final CRE clustering, along with the supervised model and important sequence features of each cluster.

#### 0.1 STOIC run

**Step 1:** An initial centroid (a CRE) is randomly chosen in the activity profile space. All CREs with an activity profile that show a correlation with the profile of the centroid above a certain threshold (initially 0.8) are included in the cluster. The centroid and the correlation threshold define the two parameters of the cluster.
**Step 2:** A supervised model (RF) is fitted to predict the probability of each CRE to belong to the cluster using only sequence features as predictors (see 0.2). The accuracy (AUC) of this model is stored, and the cluster is then optimized (the centroid and correlation radius are optimized to maximize the AUC this first model, see 0.3). If the optimized cluster has an AUC higher than a user-defined AUC threshold (0.7 in our the neuronal differentiation case study), it is added to the list of optimized clusters. After an optimized cluster is added, the coverage of the activity space (the fraction of CREs covered by a cluster) is computed. While coverage increases (it has increased by more than 10% in the last 5 added clusters), a new, distant, centroid is picked (see 0.4) and Step 2 is reiterated.
**Step 3:** Once coverage stops increasing, the list of optimized clusters is pruned to remove redundant clusters (see 0.5), and CREs are assigned to one cluster at most (see 0.6). The list of optimized clusters and supervised models is returned.

#### 0.2 Supervised model of a cluster

A probability Random Forest (RF) [Malley et al., 2012] from the ranger R package is trained to predict whether each CRE belongs to the cluster (1) or not (0), based the sequence features matrix. The default value for the number of trees is 500 trees but can be changed by the user. The minimum leaf size can also be changed, set to $0.025 * N_{CREs}$ by default. The RF estimation can be multi-threaded. The performance of the RF to predict cluster membership is measured by the Aurea Under the ROC curve (AUC) on Out-Og-Bag observations.

#### 0.3 Cluster optimization

1. **Cluster radius optimization: the cluster size is adjusted to improve the AUC.** A series of candidate correlation thresholds are tested, and if one of them improves the AUC (without model re-training, to speed to process), it is selected to be tested for a RF refit. The 8 candidate correlations are spread between the current correlation threshold $\pm 20\%$. The first correlation threshold to improve the AUC after re-training is selected to replace the former correlation threshold. The new model and its AUC replace the current cluster model and AUC.

2. **Cluster centroid optimization: the cluster centroid is adjusted to improve the AUC.** A series of different candidate centroids are tested and if one of them improves the AUC (without model re-training, to speed to process), it is selected to be tested for a refit. The candidate centroids are all CREs contained within the current cluster (for speed, if the current cluster is bigger than 500 CREs, only the 500 CREs the most correlated to the current centroid are selected). The first candidate centroid to improve the AUC after re-training is selected to replace the former centroid. The new model and its AUC replace the current cluster model and AUC. The number of model re-fits is limited to 10 for the sake of speed.

These two steps are repeated until none of the candidate centroids can improve the current cluster AUC, which stops the optimisation procedure are returns the cluster and its model.

#### 0.4 Picking a new centroid

To explore the activity space efficiently, when a new cluster is created, its initial centroid is chosen as the most distant CREs from existing centroids (the centroids of optimized clusters and the initial centroids of other clusters). This allows to start optimizing a new cluster from a region where no other cluster has already been initialized, or where no other cluster has converged. To do so, the correlation to the closest (most correlated) existing centroids in computed for each CREs, and the CRE with the smallest correlation to its closets existing centroid is chosen as the new centroid.

#### 0.5 Cluster pruning

Several clusters can sometimes start with different initial centroids but converge to the same area of the activity space during their optimization, and exhibit similar sequence features. In order to get a set of final, non-redundant clusters, they are pruned based on several characteristics. Lets define $C_i$ as the set of CREs belonging to cluster $i$. For a pair of clusters $(C_1, C_2)$, the three characteristics used for pruning are:

- their **overlap** $O_{12}$, their amount of shared CREs.

$$O_{12} = \mathrm{Jaccard(C_1, C_2)} = \frac{C_1 \cap C_2}{C_1 \cup C_2}$$

- the **correlation** of their activity profile $C_{12}$.

$$C_{12} = \mathrm{Cor}(\overline{\mathrm{profile}_1}, \overline{\mathrm{profile}_2})$$

  with $\overline{\mathrm{profile}_i}$ the vector of size $N_{samples}$ of average expression profile of the CREs in $C_i$.

- their **swap** score $S_{12}$, the similarity of the sequence rules learned by their supervised model.

$$S_{12} = \min(\mathrm{swap}_{12}, \mathrm{swap}_{21})$$

  with

$$\mathrm{swap}_{ij} = \frac{AUC_{ij}}{AUC_j}$$

  and $AUC_{ij}$ the AUC achieved when using the model learned on cluster $i$ for predicting the classes formed by cluster $j$, and $AUC_j$ the AUC achieved when using the model learned on cluster $j$ for predicting cluster $j$.

The condition for $C_1$ and $C_2$ to be considered redundant is:

$$O_{12} > \frac{2}{3} \quad \vee \quad C_{12} > 0.95 \quad \vee \quad S_{12} > s \quad \wedge \quad C_{12} > c \quad \wedge \quad O_{12} > 0$$

with $s$ a swap score defined by the user and set to 0.95 by default, and $c$ the maximum of the two final correlation thresholds obtained during optimization. When two clusters are considered redundant, only the one with the best AUC is selected.

#### 0.6 Assigning CREs to one cluster at most

It can happen that two clusters overlap (share CRE(s)), which implicates that some CREs belong to more than one cluster at the same time. In order to return a partition where each CRE belongs to at most one cluster, we rely on the CRE posterior probability of belonging to each cluster, which corresponds to the predicted value of the probability Random Forest Malley2012. A CRE is assigned to the cluster where it ranks higher in terms of posterior probability, meaning that it joins the most likely cluster given its sequence features.

#### 0.7 Feature importance measure

Two options are available in STOIC to compute feature importance:

- **Permutation importance**, measured by the drop in prediction performance caused by shuffling a feature $X_j$ on Out-Of-Bag observations as originally proposed by L. Breiman when introducing Random Forests [Breiman, 2001]. This is implemented in the ranger package, through the `importance = 'permutation'` argument.

- **Conditional permutation importance**, an extension of permutation importance with the objective of reducing bias caused by correlation between features [Strobl et al., 2008]. In this approach, the shuffling of Out-Of-Bag observations is performed within partitions of the values of the predictors in each tree that are correlated with $X_j$ over a certain threshold. This strategy is implemented in the fonction `randomForest` of the extendedForest R package, launched in STOIC with arguments `corr.threshold = 0.5, maxLevel = 8` (the latter corresponds to the level to consider for partitioning feature values in trees).

Feature importance is computed only on the final set of pruned, optimized models. The rest of the Random Forest fits made during clusters optimizations are run with permutation importance disabled (`importance = 'impurity'` in ranger, to speed up computations).

### B. Detailed experimental protocols

The following sections details the full experimental protocols for cell culture, treatment, extraction, sequencing, and bio-informatic pre-processing for the different datasets analyzed in the article.

#### Cell models and differentiation

The cell model and the complete differentiation protocol from iPSC to NSC to cortical neuron were described previously [Yip et al., 2024]. Human iPSCs line i3N is a gift from Dr. Michael Ward from NIH and derived from the WTC11 iPS by introducing a doxycycline-inducible mNGN2 transgene at the AAVS1 site (Wang et al. 2017). The iPSCs were cultured at a density of 0.5 million cells on a iMatrix-511 (Nippi; Cat.No. 892012)-coated 10 cm-dish in StemFit medium (Ajinomoto; Cat. No. RCAK02N) with 10 nM Rho-associated kinase (ROCK) inhibitor (Y-27632, Wako; Cat.No. 036-24023). The following day, ROCK inhibitor was withdrawn by refreshing the StemFit medium. After culturing for 4 days, the cells were washed with DPBS and detached by incubating with Accutase (Sigma; Cat.No. A6964) at 37 °C for 10 min. NSCs were differentiated and maintained by using the PSC Neural Induction Medium (NIM) (Gibco; Cat.No. A1647801) according to the manufacturer's instructions. iPSCs were seeded onto an iMatrix511-coated 6-well plate with 10 nM ROCK inhibitor. The following day, the medium was replaced with 2.5 mL of NIM. On day 6 of neural induction, the NSCs (P0) were harvested by incubating with Accutase at 37 °C for 10 min and scraping. The cells was replated to a iMatrix511-coated 10-cm dish with PSC Neural Expansion Medium (NEM) containing 10 nM ROCK inhibitor. The following day, NEM was refreshed to remove the ROCK inhibitor. To prepare the cells for transcriptomic profiling, the cryopreserved P1-NSCs were thawed and cultured in NEM with 10 nM ROCK inhibitor on an iMatrix511-coated plate. After two passages, P3-NSCs were harvested by incubating with Accutase and scraping, and subjected to single cell sequencing protocols. The differentiation of cortical neurons from NSC using doxycycline-inducible mNGN2 was performed. Briefly, the cryopreserved P3-NSCs described above were replated onto 0.1mg/mL Poly-L-ornithine (Sigma, Cat.No. P3655)-coated 10cm-dish at a density of 5 × 106 cells per dish in the differentiation medium I (50% DMEM/F12 (Gibco, Cat.No. 10565018) and 50% Neurobasal Medium (Gibco; Cat.No. 21103049) as the base, supplemented with 0.5 × Non-Essential Amino Acids (NEAA, Gibco; Cat.No. 11140050), 0.5 × GlutaMAX (Gibco; Cat. No.35050-061), 0.5 × N-2 Supplement (Gibco; Cat. No. 17502048), 0.5 × B-27 Supplement (Gibco; Cat. No. 17504-044), 2.5 $\mu$g/mL human Insulin (Sigma; Cat.No. 19278), 2 $\mu$M DAPT (Wako; Cat.No. 043-33581), 50 $\mu$M 2-Mercaptoethanol (Gibco; Cat. No. 21985023), 2 $\mu$g/mL doxycycline hydrochloride (Sigma; Cat.No. D-9891), 5 $\mu$g/mL Mouse Laminin, and 10 nM ROCK inhibitor). The following day, the same medium without ROCK inhibitor was used to replace the medium. On day 3, the medium was changed to differentiation medium II (50% DMEM/F12 and 50% Neurobasal Medium as the base, supplemented with 0.5 × Non-Essential Amino Acids, 0.5 × GlutaMAX, 0.5 × N-2 Supplement, 0.5 × B-27 Supplement, 2.5 $\mu$g/mL human Insulin, 10 $\mu$M DAPT, 50 $\mu$M 2-Mercaptoethanol, 2 $\mu$g/mL doxycycline hydrochloride, 0.5 $\mu$g/mL Mouse Laminin, 10 ng/ml brain-derived neurotrophic factor (BDNF, PeproTech; Cat.No. 450-02), 10 ng/ml glial cell-derived neurotrophic factor (GDNF, PeproTech; Cat.No. 450-10), 10 ng/ml neurotrophin-3 (NT-3, PeproTech; Cat.No. 450-03), and 0.5 $\mu$g/ml laminin). On Day 6 onwards, half of the medium was exchanged every 3 days without Doxycycline. On day 10, the differentiated neurons were incubated with dissociation buffer (50% Accutase, 50% DPBS, and $\geq$ 50 units papain (Worthington; Cat.No. LK003178)) at 37 °C for 20 min. Then, wash buffer (DMEM/F12 supplemented with GlutaMax, 10 nM ROCK inhibitor, and $\geq$ 500 Kunitz units DNaseI (Worthington, Cat.No. LK003172)) was added to the cells. The cells were collected, washed 3 times with 0.1% BSA in DPBS, and centrifuged at 150 × g for 10 min to remove the dead cell debris. The cells were subjected to 10x single-cell sequencing protocol.

Samples for single cell 5'end RNA-Seq were extracted using the Chromium Next GEM Single Cell 5' Library and Gel Bead Kit v1.1 (PN: 1000165). Samples for single nuclei ATAC-Seq were extracted using the Chromium Next GEM Single cell ATAC Kit v1.1 (PN: 1000175).

#### CUT&Tag dataset

Briefly, iPSCs, NSCs, and cortical neurons were harvested and subjected to nuclei isolation according to the 10x demonstrated protocol (Document CG000169 Rev D), with the concentration of BSA in PBS optimized to 0.1%. The isolated nuclei from two biological replicates were used for library preparation by CUT&Tag-IT Assay Kit (Active motif, Cat.No. 53160), according to the manufacturer's instructions. Briefly, the nuclei were immobilized to Concanavalin A coated beads, followed by magnetic separation to isolate the nuclei. The nuclei were then resuspended with antibody buffer containing protease inhibitor cocktail and digitonin. Then, 1 $\mu$g specific antibody (CTCF, H3K4me3, H3K27me3, H3K4me1, and H3K27ac) or 1 $\mu$g rabbit IgG isotype control antibody was added to each sample. The nuclei were isolated by magnetic separation after overnight incubation. The samples were then incubated with 1:100 diluted guinea pig anti-rabbit secondary antibody in Dig-Wash buffer for 1 hour at room temperature, followed by washes with Dig-Wash buffer. The antibody-bound chromatin was incubated with 1:100 diluted CUT&Tag-IT Assembled pA-Tn5 transposomes in Dig-300 buffer for 1 hr at room temperature, followed by stringent washes with Dig-300 buffer. The antibody-bound chromatin was subsequently sheared, and sequence adaptors were added by tagmentation for 1 hour at 37°C. The samples were de-crosslinked with buffers containing 0.1% SDS, 16mM EDTA, and 1 $\mu$g Proteinase K for 1 hr at 55°C, followed by DNA extraction. The libraries were amplified by PCR with index primers, and size selection was performed with SPRI beads to remove primer dimers. Sequencing was perfomed by HiSeqX_Ten (illumina) with the following conditions: R1: 150 cycles; R2: 150 cycles; Index1: 8 cycles; Index2: 8 cycles.

The raw sequencing data from CUT&Tag was processed by the ENCODE ATAC-seq pipeline (v1.7.0). Briefly, the resulting reads were mapped to hg38 human genome using Bowtie2 (v2.2.6). After removing PCR duplicates, bam files of the two replicates for each cell type were subjected to MACS2 (v2.1.0) for peak calling. The significant peaks ($p<0.01$) were merged and extended into a minimum window size of 150bp.

Chromatin states of CREs were defined by ChromHMM [Ernst and Kellis, 2012] based on 200-bp bins using the CUT&Tag and scATAC-seq data divided into 3 cell-types. This resulted in 16 clusters that were manually annotated according to their chromatin features, and to the coordinates of the SCAFE TSS clusters and SCREEN cCREs indicating promoter, enhancer and CTCF sites. The derived chromatin states, promoter (promoter and promoter flanked), bivalent promoter, active enhancer, repressed enhancer, primed enhancer, and CTCF-alone were then intersected with the CRE regions in each cell-type. If more than one state overlapped with a CRE, a representative state was chosen based on the order of chromatin states stated above. Finally, by combining chromatin states in the 3 cell types, CREs were classified as "promoter-like", "enhancer-like", "CTCF-alone" or "unclassed".

#### ChIP-Seq datasets

The iPSCs, NSC, and cortical neuron cells were harvested as described above. The cell fixation and chromatin shearing were performed by truChIP Chromatin Shearing Kit (Covaris, Cat.No. 520127) according to the manufacturer's instructions with some modifications. Briefly, 3 × 107 cells were crosslinked with 1% methanol-free formaldehyde (Electron Microscopy Sciences) in PBS for 5 min at room temperature, followed by quenching with 200 mM glycine for 5 min. The fixed cells were lysed, and the chromatin was sheared using the Covaris S220 ultrasonicator. To validate the chromatin shearing efficiency, the sheared chromatin was de-crosslinked in 50 mM NaCl for 20 min at 100°C, followed by ramping the temperature down to 50°C. 500 ng of DNA was examined by 1.5% agarose gel electrophoresis. After confirming the chromatin shearing efficiency within the range of 200-1200 bp, the chromatin immunoprecipitation reaction was performed using the ChIP-IT High sensitivity (Active motif, Cat.No. 53040) with Spike-in Antibody (Active motif, Cat.No. 61686) and Spike-in chromatin (Active motif, Cat.No. 53083) according to the manufacturer's instructions. Sequence libraries were prepared using the Next Gen DNA Library Kit (Active Motif, Cat.No. 53216). The resulting libraries were quantified by Qubit 3.0 Fluorometer using Qubit dsDNA HS Assay Kit (Invitrogen, Cat.No. Q32851) and were paired-end sequenced by NovaSeqX plus (illumina) with 150 bp read length and 8 bp single-indexed reads. The fastq files were mapped to the hg38 genome with BWA (Li and Durbin 2010). The bam files were porcessed with fixmate and markdup by samtools (Danecek et al. 2021) while the bam files free of duplicates were applied to peak calling. Peak calling was performed by MACS2 (Zhang et al. 2008) at FDR 0.01 for each replicate using the pair-end bam files from specific and isotypic antibodies. The peaks of the 2 replicates were merged by bedtools as a permissive output, and were subjected to IDR (Li et al. 2011) for reproducible peaks at p-value $< 0.05$.

#### Hi-C dataset

After harvesting the cells (iPSC, NSC and Neuron), 2 replicates of each cell type were fixed by 1% formaldehyde and Hi-C library construction was performed with the Arima-HiC+ kit and the Arima Library Prep Module (Arima Genomics) according to the manufacturer's protocols. The libraries were then subjected to Illumina NovaSeq6000 for sequencing with 150 bp paired-end mode. We first trimmed the 6 bp from the left end of each read as suggested by Arima Hi-C data analysis and then trimmed the 70 bp from the right end of each read using the command line "seqtk trimfq -b 6 -e 70". The remaining 75 bp reads were used for the nf-core hic pipeline v 2.1.0 (https://nf-co.re/hic/2.1.0/). The two replicates from the same cell type were pooled together for the final nf-core Hi-C pipeline.
